# Resource availability and dimensionality result in ecology-dependent selection in bacteriophage spatial expansions

**DOI:** 10.64898/2026.02.23.707387

**Authors:** Hassan Alam, Diana Fusco

## Abstract

In microbial populations, fitness, which is essential to understand and predict evolution, is often defined and measured as the net growth rate of a population in isolation. Applying the same definition to viruses is challenging, both because viral replication involves a host infection process, which is determined by several parameters that are context-dependent, and because viruses compete heavily for resources (susceptible cells). These challenges are particularly exacerbated in spatial range expansions, where multiplicity of infection is often high and resource availability varies in time and space.

To assess different fitness definitions and their generalizability, we investigate a model of coupled partial differential equations for phage plaque expansion in one and two dimensions. We find that two commonly used metrics for phage fitness in plaque expansions, i.e., steady state phage densities and front expansion speed in isolation, are unable to reliably predict the winner in one- and two-dimensional direct competitions. More generally, we find that optimal phage traits depend on the dimensionality of the system and the make-up of the phage population, leading to unexpected behaviours, e.g., rock-paper-scissor dynamics and, in high dimensions, enhanced phage density due to the nearby presence of a competitor. We show that the phenomenon stems from the interplay between resource consumption and replication and thus may apply more broadly to any population competing for shared resources.

## 1. Introduction

Bacteriophages, also called phages, are viruses that infect and replicate within a host bacterium and play a critical role in microbial communities in the wild, shaping their ecology, structure, and evolutionary dynamics [1–6]. Virulent phages, in particular, which rely on host lysis for propagation, have recently received renewed interest because of their theurapeutic potential against antibiotic resistant bacteria [7–9].

Given their large numbers, phages in nature often experience high levels of competition for bacterial resources [10–13]. While ecological niches are known to occur, so that phages specialize in infecting specific hosts [14], it is still estimated that on average each bacterial strain can be infected by at least 10 distinct phages [15–17]. To avoid resource sharing and limit competition, phages have evolved a broad array of mechanisms to prevent superinfection (the phenomenon by which a phage infects a previously infected host) [18–23], whose fitness (dis)advantages we have recently investigated in simulations of well-mixed cultures [24]. Despite the ubiquitousness of phage and their well-established role in microbial evolution, a clear and broadly applicable definition of phage *fitness* that accounts for the inevitable resource competition they experience is still missing.

In the literature, different definitions of phage fitness have been used depending on the research question and the environment. For example, in the case of phage plaque expansion within a bacterial lawn, plaque size, plaque front expansion speed, phage growth rate, total number of phages and phage density inside the plaque have all been used as a measure of phage fitness, both in experiments and theoretical work [25–36]. All these measures, however, assess the properties of a single phage population growing in isolation, which have been shown to be poor predictors of the outcome of phage competitions [24, 37]. Alternative measures, such as the ratio of competing phage populations inside a plaque, might be more appropriate to quantify phage relative and competitive fitness as they take into account the competition for shared resources between the two viruses [33], however, the predictive power of these different observables and their transferability from one system to another have not been adequately explored [38]. For instance, in a two dimensional spatial range expansion with two phage populations competing for a single resource (one bacterial strain), the ability of phage to occupy uninfected space at the plaque front is expected to be a more appropriate measure of phage fitness than phage density alone [39–44]. Taking all these observations together, we currently do not know whether a phage that is theoretically predicted to emerge in an evolutionary experiment would actually establish in an experiment and whether the selective pressures predicted in some conditions are truly present. Reciprocally, a poorly defined fitness metric makes it difficult to pinpoint the reason why specific phage populations outcompete others both *in vitro* and *in vivo*.

To address this gap, here we extensively study two different definitions of phage fitness that have been routinely used in the literature to predict phage evolution in plaque expansions: (i) maximum steady-state phage density at the plaque front, and (ii) steady-state plaque front expansion speed. By computing these quantities as a function of lysis time and adsorption rate in phage populations grown in isolation, we determine their predictive power both in 1D- and 2D-expansion competitions. Interestingly, despite the simplicity of the setup we study, not only we find that both measures in isolation are poor predictors of the outcome of a direct phage competition, but also that the competition outcome depends on the dimensionality of the system. Careful investigation of the results shows that this observation originates from the fact that phage growth dynamic is deeply dependent on the surrounding coexisting phage populations because of resource limitations, to the point that a mixture of two phages expanding at the same steady-state speed in isolation display a significant slow-down when combined. This result leads to interesting and unexpected behaviour, such as potential rock-scissor-paper dynamics, where the optimal phage traits under selection depend on the make-up of the phage population. Our results highlight not only overlooked problems in applying a single phage fitness definition across different contexts, but also, more intriguingly, the remarkable properties of phage fitness, which cannot be easily deconvolved from the corresponding ecological dynamics.

## 2. METHOD

To simulate phage competition, we employ a system of non-linear spatiotemporal reaction-diffusion equations that captures the population dynamics of phage-bacteria systems in one- and two-dimensions. As in the classic model system coliphage T7, we assume post-adsorption superinfection exclusion [45–48] in which superinfecting phages are not allowed to replicate after their adsorption onto the cell wall and they effectively die. The model can accommodate multiple potential phases during the infection cycle, e.g., viral DNA injection, viral DNA replication, virion assembly, etc. (figure 1a). However, to keep the number of parameters small, we use only two phases in the following analysis corresponding to the eclipse and the productive period of the infection cycle. This implementation leads to a cell lysis time that is not deterministic or exponentially distributed as in majority of previous work [49–52], but follows a hypoexponential distribution [53], which is more representative of experimental lysis time distributions that display a well-defined mode with some degree of variation around it [54].

**Figure 1.**
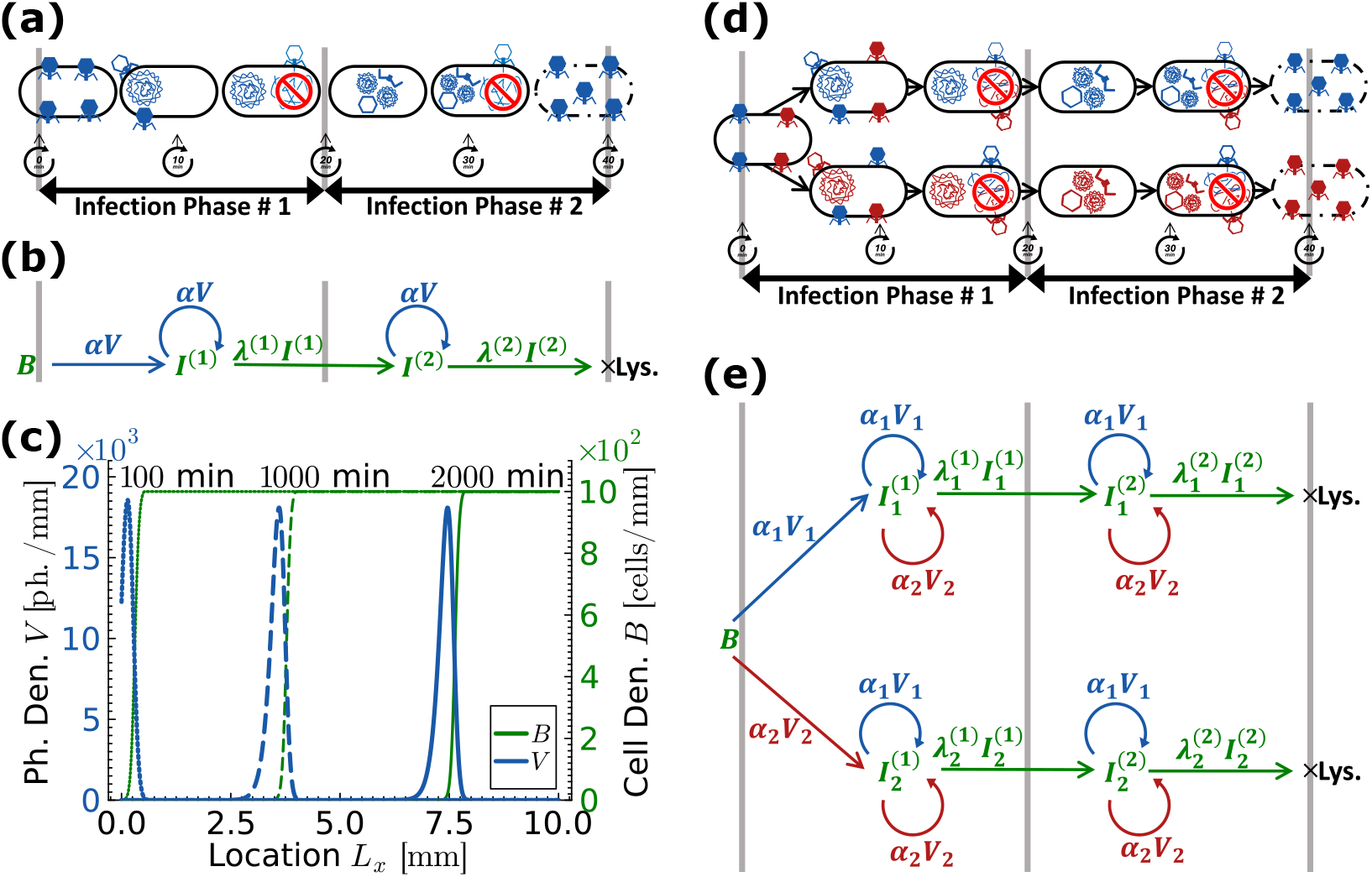
Schematics, infection flow, and population dynamics plots of one phage and two phage strains models with superinfection. (a) Schematic of the one phage and one bacterial strain model. It shows that the first phase of infection starts when a phage attaches to a bacterial cell. The phage then injects its DNA into the cell. During this phase, the infected cell may be superinfected by more phages. However, superinfecting phages are not allowed to replicate in the model. The infection continues to the second phase during which progeny phage components assemble inside the cell. During this phase, the infected cell may again be superinfected by more phages. Finally, the infected cell lyse and progeny phages are released. (b) Infection flow diagram of the one phage and one bacterial strain model. In the diagram, *V* is the density of phages, *B* is the density of uninfected bacterial cells, and *I*^(*k*)^ is the density of infected bacterial cells that are in the *k*^*th*^ phase of infection, where *k* = 1, 2. *α* is the adsorption rate of the phages. *λ*^(*k*)^ are the infection progression rates of infected cells *I*^(*k*)^. (c) Population dynamics of the one-dimensional one phage and one bacterial strain model. The figure shows population density distributions of phage and bacteria strains in the system at time points 100, 1000, and 2000 minutes. It shows that the plaque is expanding in the system from left to right with time. Bacterial cell density is shown by the green curve and phage particle density is shown by the blue curve. Adsorption rate is taken as 5 × 10^−6^ mm*/*cell · min, and all remaining parameters are mentioned in table 1. (d) Schematic of the two phage and one bacterial strain model. It shows that the bacterial cells may be infected by either the blue phage or the red phage strain. (e) Infection flow diagram of the two phage and one bacterial strain model. In the diagram, subscripts denote the type of the infecting phage — 1 denotes the blue phage and 2 denotes the red phage strain.

Figure 1 shows a schematic of the infection cycle and the population dynamics in our model for the case of one phage strain vs. one bacteria strain (a-c) and two phage strains vs. one bacteria strain (d-e). First, an uninfected bacterial cell is infected by a phage marking the beginning of the first phase of infection. The infection progresses at a rate *λ*^(1)^ to the second phase, and then at a rate *λ*^(2)^ to lysis. The infected cell may be superinfected by secondary phages at any point during the infection. However, superinfecting phages are not allowed to replicate and are lost without changing the fate of the infected cell. Table 1 summarizes the model parameters, and the initial and boundary conditions used throughout the study.

**Table 1.** Model parameter values and units used in the study, except stated otherwise.

| One-Dimensional System<sup>(a)</sup> |  |  |  |
| --- | --- | --- | --- |
| Parameter | Symbol | Unit | Values |
| Initial Cell Density | $B$ | cells/mm | 1000<sup>(b)(c)</sup> |
| Initial Phage Density | $V$ | phages/mm | $N(\mu = 0.0, \sigma = 0.01)$<sup>(d)</sup> |
| Initial Total No. of Phage | — | phages | 150 |
| Finite Difference Step Size | $\Delta x$ | mm | 1/150 |
| Discreteness Threshold | $\theta = 1/\Delta x$ | ph./mm | 150 |
| Hill Coefficient | $M$ | — | 10 |
| Phage Diffusion Rate | $D$ | mm<sup>2</sup>/min | 0.00024 |
| Phage Decay Rate | $\delta$ | min<sup>-1</sup> | 0.05 |
| Burst Size | $\beta$ | phages | 100 |
| Adsorption Rate | $\alpha$ | $\frac{\text{ph.} \cdot \text{mm}^{-1} \cdot \text{min}^{-1}}{\text{cells} \cdot \text{mm}^{-1} \cdot \text{ph.} \cdot \text{mm}^{-1}}$ | 0.01 |
| First Phase Infec. Progr. Rate | $\lambda^{(1)}$ | min<sup>-1</sup> | 0.05 |
| Second Phase Infec. Progr. Rate | $\lambda^{(2)}$ | min<sup>-1</sup> | 0.05<sup>(e)</sup> |

<sup>(a)</sup> **For two-dimensional system:** The step size is $\Delta x = \Delta y = 1/150$ mm. Discreteness threshold is $\theta = 1/\Delta x \Delta y = 150$ ph./mm<sup>2</sup>. Hill coefficient is $M = 10$ . Unit of cell density is cells/mm<sup>2</sup>, phage density is phages/mm<sup>2</sup>, and adsorption rate is $\frac{\text{ph.} \cdot \text{mm}^{-2} \cdot \text{min}^{-1}}{\text{cells} \cdot \text{mm}^{-2} \cdot \text{ph.} \cdot \text{mm}^{-2}}$ .
<sup>(b)</sup> Initially cells are uniformly distributed in the system.
<sup>(c)</sup> Relevant dynamics in our simulations are independent of the absolute values of phage and cell densities. We can rescale these densities by any factor $1/k$ (*i.e.*, $\tilde{B} = B/k$ and $\tilde{V} = V/k$ ) while simultaneously rescaling the adsorption rate by the reciprocal factor (*i.e.*, $\tilde{\alpha} = k\alpha$ ) to maintain the product $\tilde{\alpha}\tilde{B} = \alpha B$ invariant. These adjustments ensure that the relevant dynamics of the system remain unchanged.
<sup>(d)</sup> Initial phage density follows a normal distribution with a mean of $\mu = 0$ and a standard deviation of $\sigma = 0.01$ . This distribution is sharply peaked, resembling a delta function, with 150 phages located at the left boundary of the system. Open boundary conditions are applied at both boundaries of the system.
<sup>(e)</sup> Mean lysis time can be calculated by the formula $\langle \tau \rangle = \frac{1}{\lambda^{(1)}} + \frac{1}{\lambda^{(2)}}$ following the Hypoexponential distribution.

The equations for the one phage and one bacterial strain model are given below, while the corresponding equations of the two phage and one bacteria strain model are given in supplementary information.

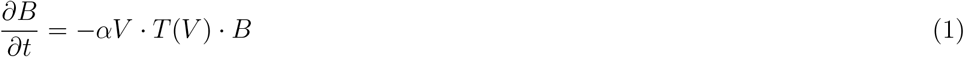

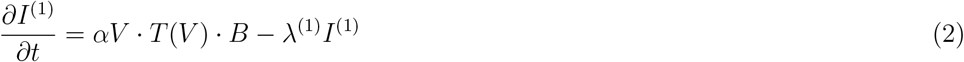

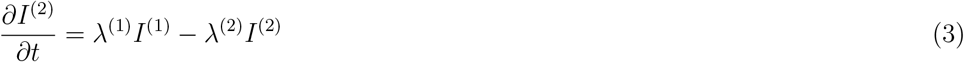

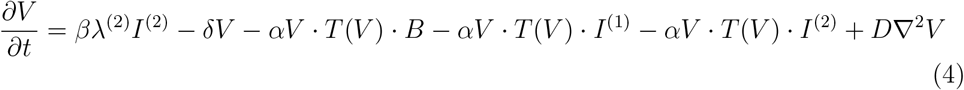

Here, *B, I*^(1)^, *I*^(2)^, and *V* are the densities of uninfected cells, cells in the first and second phase of infection, and free phages, respectively. *α* is the adsorption rate, *β* is the burst size, *δ* is the decay rate, and *D* is the diffusion rate of phages. *λ*^(1)^ and *λ*^(2)^ are infection progression rates of cells in the first and second phases of infection. Analogously to previous models of bacterial lawns, we assume that bacteria are non-motile and their growth is negligible [33, 55].

The factor *T* (*V*) is a Hill function that accounts for the discreteness of phages, by effectively removing the reaction terms that involve phage when phage density is below a threshold without introducing discontinuities in the system of PDEs [44]:

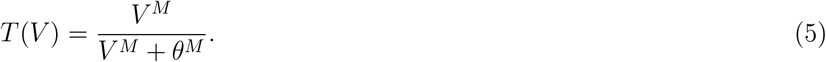

Here, *θ* is the **half-saturation constant** at which *T* (*V*) = 0.5 and corresponds to the discreteness threshold (minimum number of phages per lattice site) while *M* is the Hill coefficient. If at least one phage is needed at each lattice site to trigger the reaction terms, *θ* is equal to the inverse of the step size used to numerically integrate the partial differential equations. In the limit of large *M, T* (*V*) becomes a unit step function where the step is located at *V* = *θ*, meaning that phages can infect a bacterial cell only when *V > θ*. In this study, we have set *M* = 10 and *θ* = 150 ph.*/*mm. As a result, phage reaction terms sharply vary from 0 when phage densities are too low (less than one phage per lattice site) to finite values.

In the supplementary information, two additional models are also discussed, which are helpful to interpret some of the results in the later sections. One model does not allow for superinfection (no adsorption occurs to infected cells), while the other model has no phage discreteness threshold.

### 2.1. Definitions of phage fitness

The reproductive potential of a phage population in a microbial environment is called **phage fitness *F* (*t*)**. Its mathematical definition depends on the type of microbial system and the context of the study. For example:

(I) In steady state well-mixed liquid cultures consisting of only one phage and one bacterial strain, the steady state phage concentration has been used as proxy of phage fitness [56].

(II) In well-mixed liquid cultures that do not attain steady state, the growth rate *r*(*t*) of the phage at a time *t* has been taken as a measure of phage fitness *F* (*t*) [32, 34]. We note here that if the phage growth is exponential, then its growth rate is equal to the rate of change of the logarithm of its population *V* (*t*) over time. This can be shown by considering the following exponential growth rate equation,

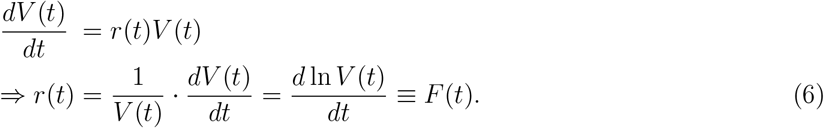

(III) Extending scenario (II) to two phage strains with populations *V*_1_ and *V*_2_ competing for a single resource (one bacterial strain) in a well-mixed liquid culture, the difference between their growth rates, *r*_1_(*t*) and *r*_2_(*t*), can be taken as a measure of their [38, 57, 58] relative phage fitness Δ*F* (*t*) as follows,

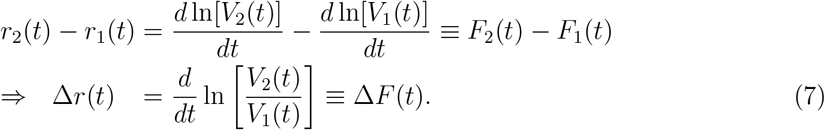

(IV) For a single phage strain replicating on a uniform bacterial lawn on a Petri dish in a spatial range expansion, then the size (radius) of the plaque, the total number of phages inside the plaque, the maximum phage density at the plaque front or the plaque front expansion speed have all been used as proxies for phage fitness [33, 59–62].

(V) Extending scenarion (IV) to two phage populations competing for a single resource (one bacterial strain) in one dimensional spatial range expansion, the definition of phage fitness given in the equation 7 can in principle be applied by considering *V*_1_(*t*) and *V*_2_(*t*) as either the maximum phage densities at the plaque front or total number of phages inside the plaque.

(VI) If two phage populations compete for a single resource (one bacterial strain) in a two dimensional spatial range expansion, then the ability of phage to occupy uninfected space at the expanding edge, like plaque front speeds, measures the phage fitness [39–44].

In this work, we investigate the following three microbial configurations to assess the transferability of distinct fitness definitions across different scenarios and their power in the predictive the outcome of direct competitions:

i. One phage and one host strain growing in one dimension.
ii. Two phages competing for one host strain growing in one dimension.
iii. Two phages competing for one host strain growing in two dimensions.

## 3. RESULTS

### 3.1. Comparison of phage fitness definitions in single-phage one-dimensional plaque expansion

We numerically solve the equations of the one-dimensional single phage strain model with superinfection and discreteness threshold using the parameters, initial conditions, and boundary conditions specified in Table 1 (further information in SI) and analyse (i) the steady state maximum phage density and (ii) the steady state plaque front speed as two distinct and often used definitions of phage fitness in spatial expansions. An example of phage and host population profiles at different time points for this microbial configuration is shown in fig. 1c, and the convergence to steady-state of the two fitness measures is shown in fig. A1. The method used to calculate the steady state values is described in the supplementary information.

Fig. 2 shows how the two fitness measures depend on phage adsorption rate, *α*, and the second phase infection progression rate, *λ*^(2)^. Both curves are dome-shaped, characterized by an optimal value of adsorption rate associated with maximal fitness, in agreement with previous work on plaque expansion [25, 33, 35, 63–65]. However, the prediction of this optimal adsorption rate differs for the two fitness measures, showing that the two definitions are not interchangeable and lead to distinct quantitative predictions.

**Figure 2.**
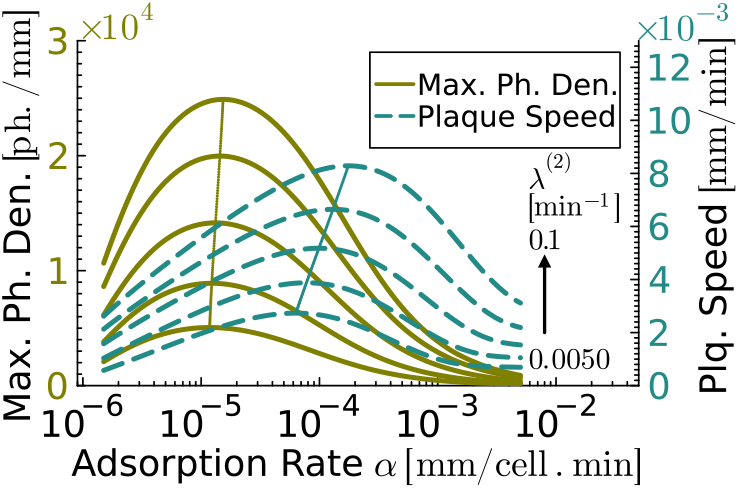
Phage fitness versus adsorption rate for the one-dimensional one phage strain model with superinfection and discreteness threshold against different values of second phase infection progression rates. Plots for two phage fitness criteria — steady state maximum phage density at plaque front (olive solid lines) and steady state plaque front expansion speed (dark cyan dash lines) — are shown in the figure at five different values of second phase infection progression rates: *λ*^(2)^ = 0.005, 0.01057, 0.02236, 0.04729, and 0.1 min^−1^ from bottom to top. The plots are dome-shaped, and lines tracing the peak of the domes are also shown on respective fitness plots. To obtain *near* steady state conditions, plaques are allowed to expand for a time of 1000 minutes when *α* ≥ 5 × 10^−6^ mm*/*cell · min and for a time of 2000 minutes when *α <* 5 × 10^−6^ mm*/*cell · min. Model parameters used to obtain the figure are mentioned in the table 1.

### 3.2. Phage fitness in isolation in unable to predict the outcome of direct competition

To test which fitness measure has higher predictive power in competition, we mapped the data from fig. 2 to isofitness curves in the (*α, λ*^(2)^) parameter space (fig. 3), we identified phages that are predicted to be neutral with respect to each other and directly competed them against each other. If two competing phage are indeed neutral, they should co-exist at the front indefinitely. If they are not, one will overtake the front over time and we can attempt to calculate its relative fitness using equation 7.

**Figure 3.**
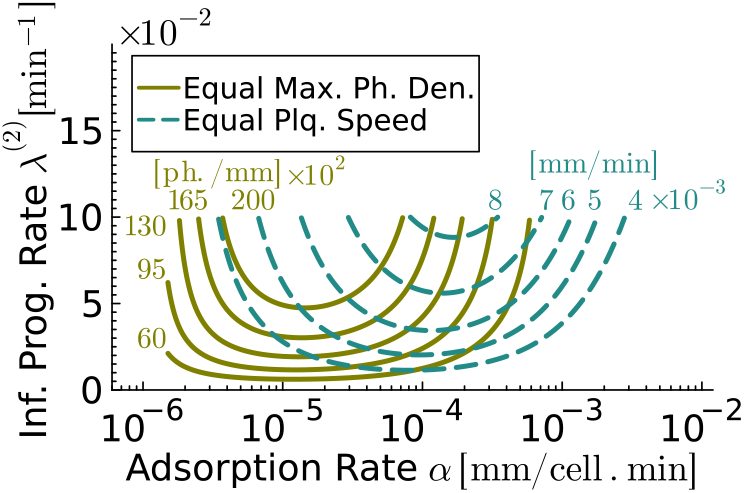
Isofitness curves for the one-dimensional one phage strain model with superinfection and discreteness threshold. The isofitness curves provide different combinations of adsorption rates and second phase infection progression rates for phages having the same fitness levels. Isofitness curves representing the steady state maximum phage densities, 6000, 9500, 13000, 16500 and 20000 ph./mm, are shown by olive solid lines and isofitness curves representing the steady state plaque front expansion speeds, 4 × 10^−3^, 5 × 10^−3^, 6 × 10^−3^, 7 × 10^−3^ and 8 × 10^−3^ mm/min, are shown by dark cyan dash lines. Model parameters used to obtain the figure are mentioned in the table 1.

We selected six phages (labelled from *A* to *F* in figures 4 and 5) for six pair-wise competitions in one-dimensional expansions. Phages *A* to *C* were selected from isofitness curves representing equal maximum phage density of 16, 500 phages/mm, while phages *D* to *F* were selected from isofitness curves representing equal plaque front speed of 5 *×* 10^−3^ mm/min. The values of adsorption rates and second infection progression rates of the selected phages are reported in table F1. Figure 4 shows that identical steady state maximum phage density in isolation fails to predict neutral pair-wise competitions — phages with higher adsorption rates consistently reach fixation in all three competitions. Similarly, figure 5 shows that identical steady state plaque speed in isolation also fails to predict the outcome of pair-wise competitions. Interestingly, in this case, the phage with the smaller *λ*^(2)^ value wins (E), and, if this is the same (competition D-F), the one with higher adsorption rate does.

**Figure 4.**
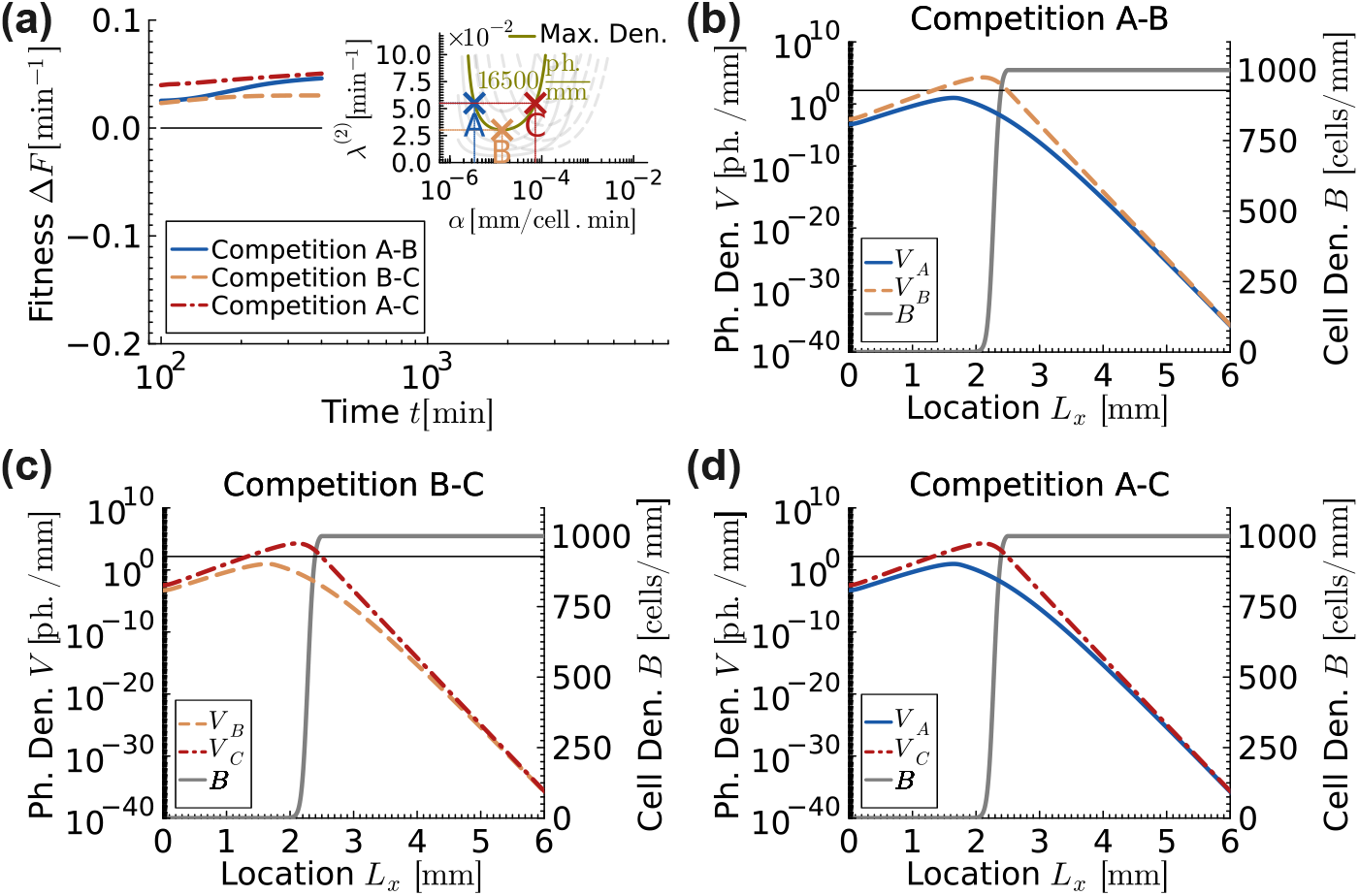
Fitness and population dynamics of phages, having equal steady state maximum phage densities of 16500 ph./mm in isolation, competing in one-dimensional uniform bacterial lawn for the one phage strain model with superinfection and discreteness threshold. (a) Relative phage fitness versus time plot with inset plot showing the three competing phages labelled as *A, B*, and *C*. Positive values of Δ*F* indicate that the second phage in the competition (B for A-B, C for B-C and C for A-C, respectively) is winning. Phages with higher adsorption rates outperform in all three cases as the fitness lines approach positive steady state values. (b) to (d) Viral and host population distributions at time 400 minutes. Gray solid lines represent the bacterial cell densities. The thin horizontal black line marks the discreteness threshold for phage density, *θ* = 150 ph.*/*mm. Below this line, phages are effectively absent. In all cases, the phage with higher adsorption rate eliminated the phage with lower adsorption rate. Adsorption rates and second phase infection progression rates of the competing phages are mentioned in the table F1, and all remaining parameters are mentioned in table 1.

**Figure 5.**
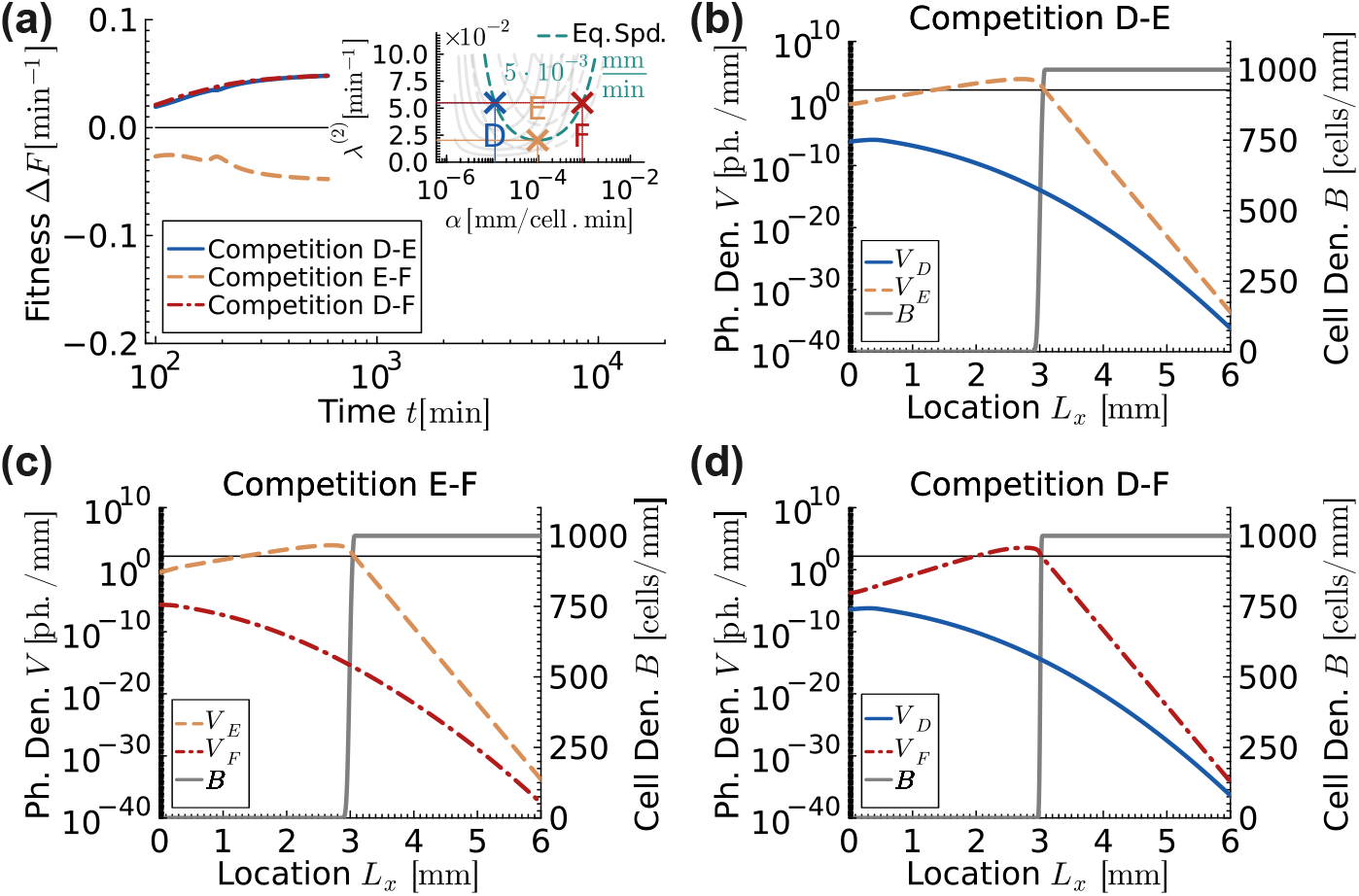
Fitness and population dynamics of phages, having equal steady state plaque front expansion speeds of 5 × 10^−3^ mm/min in isolation, competing in one-dimensional uniform bacterial lawn for the two phage strains model with superinfection and discreteness threshold. (a) Relative phage fitness versus time plot with inset plot showing the three competing phages labelled as *D, E*, and *F*. Positive values of Δ*F* indicate that the second phage in the competition (E for D-E, F for E-F and F for D-F, respectively) is winning. Phages with higher adsorption rates outperform in competitions D-E and D-F as the fitness lines approach positive steady state values. However, phage with lower adsorption rate outperform in competition E-F as the fitness lines approach negative steady state value. (b) to (d) Viral and host population distributions at time 600 minutes. Gray solid lines represent the bacterial cell densities. The thin horizontal black line marks the discreteness threshold for phage density, *θ* = 150 ph.*/*mm. Below this line, phages are effectively absent. Adsorption rates and second phase infection progression rates of the competing phages are mentioned in the table F1, and all remaining parameters are mentioned in table 1.

The observation that maximum phage density in isolation is a poor predictor of phage fitness in competition is not surprising: when competing for resources, it is paramount for a phage to get hold of a susceptible host before another phage in order to reproduce. Therefore, while a lower adsorption rate can increase phage density in isolation by enabling viral particles to diffuse further towards uninfected host, in competition, it will lower the chances for the phage to be the first to infect, effectively hindering its viability.

The reason why plaque speed in isolation fails to predict the outcome of a competition is more subtle. We found that if we removed the cutoff term in our PDEs, which accounts for the discreteness of phage particles at very low densities, plaque speed in isolation successfully predicts fitness in competition (fig. D4), pointing to the discreteness threshold as the source of the disagreement. Without the discreteness threshold, the speed of the phage population is dictated by the very front of the expansion, where density is very low and competition for resources is negligible [66–70]. When the discreteness threshold is on, the part of the expansion wave that sets the speed shifts to higher densities where competition starts playing a role, leading to a different dynamic in competition and in isolation.

Another important feature that we observe in both cases is that the value of relative fitness is not additive. This is very obvious in Fig. 5a: we see that phages E and F have the same relative fitness Δ*F* with respect to D, but the relative fitness between them is not 0 (Δ*F*_*DF*_ ≠ Δ*F*_*DE*_ + Δ*F*_*EF*_), implying that we cannot easily define an absolute fitness scale for phage competitions. The dynamics of a phage population strongly depends on the other competing phage(s), as expected when density-dependent selection is strong [71–76], which has strong implications for phage ecologies that are characterized by high degrees of diversity [77].

A consequence of the density-dependent nature of phage competition is that one cannot easily define a reference phage strain that can be used to calculate phage outcome with respect to. This is exemplified in figs. 6 and 7. If we were to use as reference phage F from fig. 5, we can in principle find the combinations of adsorption and infection progression rates that correspond to phage strains predicted to expand at the same speed as phage F in isolation (dark cyan dashed line in fig. 6) and to coexist with F in 1D competition (black solid, see also fig. 7c-d). Since both phages U and W can co-exist with phage F, one would naively expect that would be able to co-exist with each other. Direct competition (fig. 7a-b) clearly shows that U, which corresponds to the faster phage in isolation, outcompetes W.

**Figure 6.**
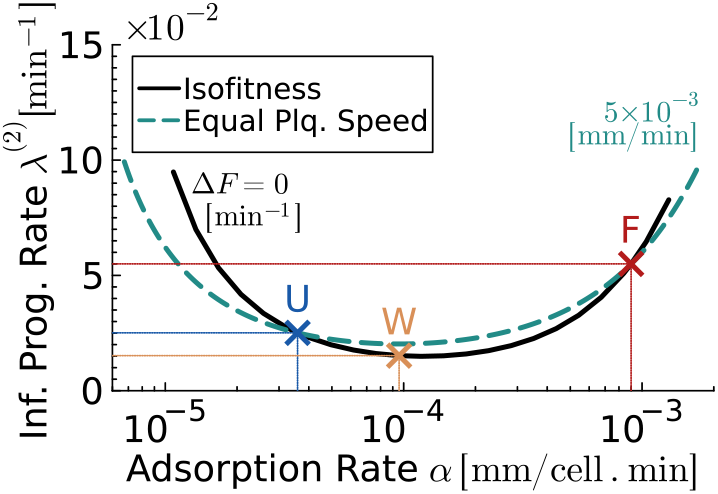
Isofitness curves for the one-dimensional one phage strain model and one-dimensional two phage strains model. Both models are with superinfection and discreteness threshold. For one phage strain model, in which phages replicate in isolation, the isofitness curve represents phages having equal steady state plaque front expansion speed of 5 × 10^−3^ mm/min (dark cyan dash line). For two phage strains model, in which phages replicate in competition, the isofitness curve represents phages having zero relative fitness Δ*F* = 0 min^−1^ relative to the phage *F* as per the definition in equation 7 (black solid line). The two intersection points, phages *U* and *F*, between the two isofitness curves represent the pair of phages that are neutral in one-dimensional competition and have the equal steady state plaque front expansion speed of 5 × 10^−3^ mm/min in isolation. Phage *W* has the same adsorption rate as phage *E* in figure 5 but the infection progression rate is reduced (slow down) to ensure that it is neural relative to the phage *F* in one-dimensional competition. Adsorption rates and second phase infection progression rates of competing phages are mentioned in the table F1, and all remaining parameters are mentioned in table 1.

**Figure 7.**
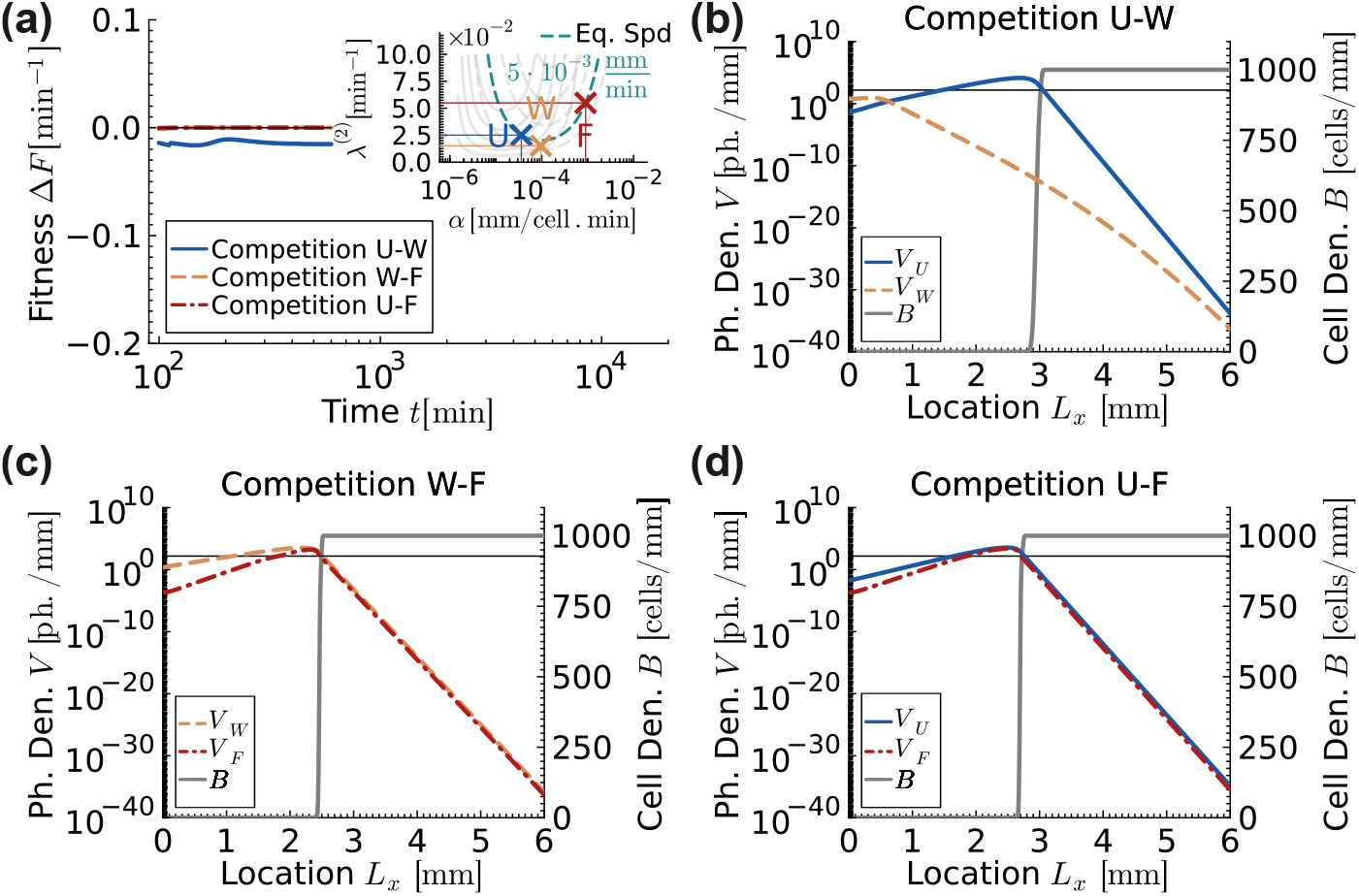
Fitness and population dynamics of phages that are neutral relative to phage *F* when competing in one-dimensional uniform bacterial lawn for the two phage strains model with superinfection and discreteness threshold. (a) Relative phage fitness versus time plot with inset plot showing the three competing phages labelled as *U, W*, and *F*. Positive values of Δ*F* indicate that the second phage in the competition (W for U-W, F for W-F and F for U-F, respectively) is winning. Phages *U* and *W* are neutral relative to phage *F* in competitions U-F and W-F as the fitness lines are at Δ*F* = 0 min^−1^ level. However, phage *U* with lower adsorption rate outperform in competition U-W as the fitness level is negative. This further demonstrates that relative fitness is not additive. (b) to Viral and host population distributions at time 600 minutes. Gray solid lines represent the bacterial cell densities. The thin horizontal black line marks the discreteness threshold for phage density, *θ* = 150 ph.*/*mm. Below this line, phages are effectively absent. Adsorption rates and second phase infection progression rates of the competing phages are mentioned in the table F1, and all remaining parameters are mentioned in table 1.

The relative position of the isofitness curves in fig. 6 also provides a clearer intuition of how phage competition affects the phage dynamics. We see that at low adsorption rates, a phage has to have higher *λ*^(2)^ (higher speed) than predicted in isolation to keep up with F, since its low adsorption rate severely reduces the chances to get a hold of a host before the competing phage. At intermediate adsorption rates, F would be outcompeted unless the competing phage slows down. This is likely because F is losing a lot of infection opportunities to superinfection in the presence of a competing phage. Indeed, as adsorption keeps increasing, phages need again to speed up to keep up with F as they waste even more phages than F to superinfection. An orthogonal way of thinking about the process is that the presence of the competing phage W slows down F. We will elaborate on this point more in the next section.

### 3.3 Dimensionality of the spatial range expansion changes competition outcome

In figures 4 and 5, we have found that phage fitness measured in isolation fails to predict the outcome of a two-phage competition and that, more generally, the density-dependent nature of phage competition prevents the definition of a relative fitness scale more in general. We now dig deeper and investigate whether the outcome of a phage competition in a one-dimensional expansion predicts the outcome of the same competition in two dimensions. Modelling efforts often reduce plaque expansions, which are in practice quasi-two-dimensional or three-dimensional depending on the experimental settings, into a one-dimensional system for simplicity and computational efficiency [30, 61, 78]. Therefore, assessing whether fitness results are independent of dimension is crucial to understand whether modelling results obtained in one dimension can be applied to higher dimensional systems, including experiments.

We initialized the two-dimensional pair-wise simulations such that the top half of the system contained only the phage with higher adsorption rate, while the bottom half of the system contained only the phage with lower adsorption rate. This side-by-side initial configuration is meant to assess phage competition at the front of the expansion as it would emerge after the unavoidable strain segregation and sectoring pattern formation that arises due to allele surfing occurring during range expansions [79–81]. To limit the transient effect of the initial condition relaxing to steady-state, we initialized the simulations with the steady-state population distribution profiles for both phages and bacteria. These profiles were obtained by first running one-dimensional simulations of the one phage model with superinfection and discreteness threshold of the respective phages up to a plaque size (radius) of at least 1000 mm.

Snapshots of three different two-dimensional pair-wise competition simulations at 900 and 1350 minutes are shown in fig. 8 and a zoomed-in front snapshots at long times (20,000 minutes) are shown in fig. 10. Because phage continues to diffuse behind the front, we track both the *instantaneous equal phage density line* (thick, black) and the *historical equal phage density line* (cyan). The first shows that (*x, y*) position where the two competing phages have the same density at the time of the snapshot. The second tracks the intersection between the equal phage density line and the maxima of the total phage density profile (expansion front) over time. Figures 9 and 10 also shows the phage density profiles along the expansion direction (*x*-coordinate) at the expansion front on the equal phage density line (cyan cross in the left column) at short and long timescales, respectively.

**Figure 8.**
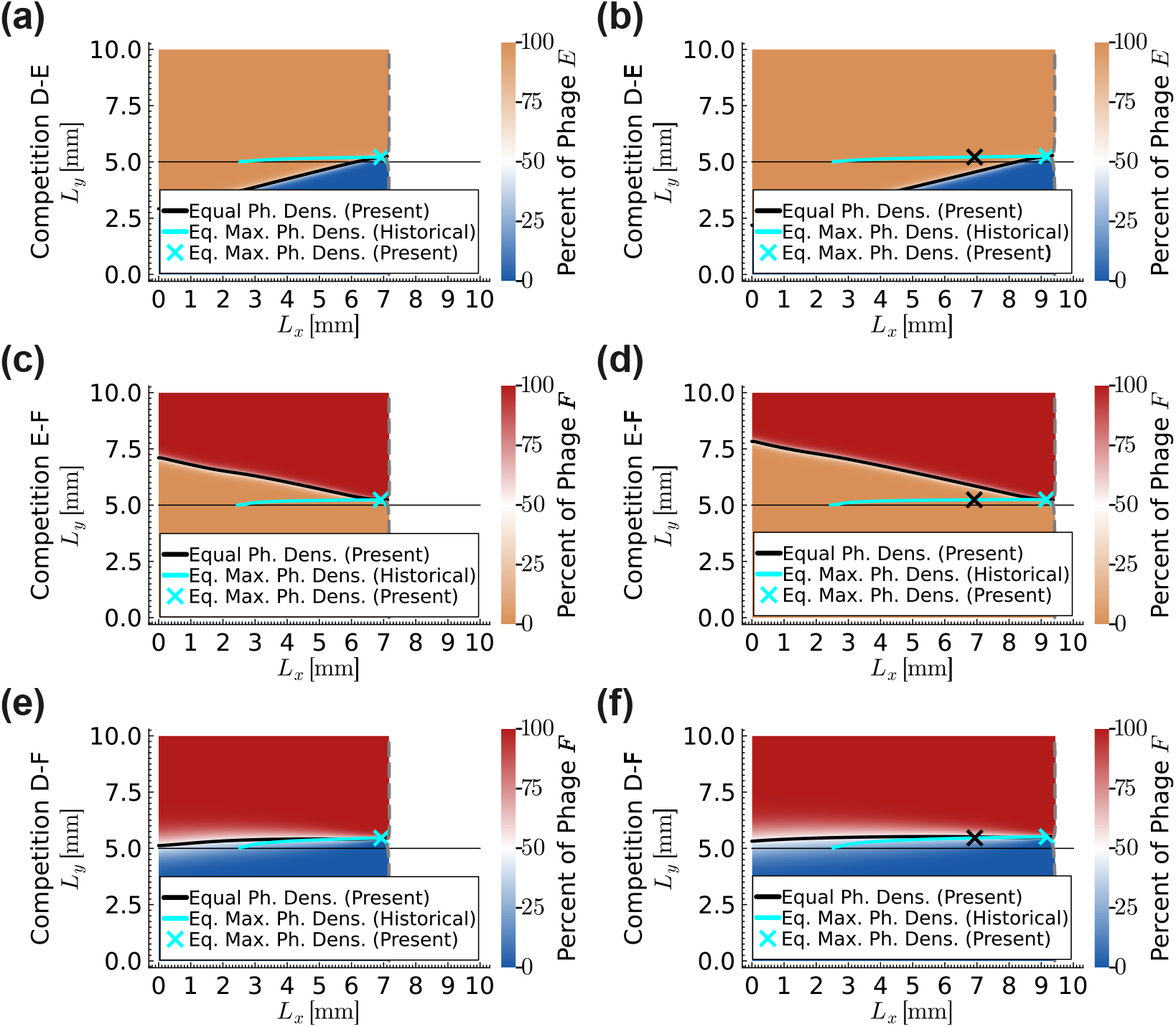
Population dynamics of phages competing in a two-dimensional uniform bacterial lawn at a short timescale for the phages with equal steady state plaque front expansion speeds of 5 × 10^−3^ mm/min in isolation for the model with superinfection and discreteness threshold. Phages *D, E* and *F*, having equal steady state plaque front expansion speeds of 5 × 10^−3^ mm/min in isolation, are competed pair-wise in two-dimensional uniform bacterial lawn. Snapshots of the simulations are given at times 900 (left column) and 1350 minutes (right column). Different phage dominated regions are shown by their respective colour labels (blue colour for phage *D*, orange colour for phage *E* and red colour for phage *F*). Vertical gray dashed line is the plaque front. Black solid line is the instantaneous equal phage density line and cyan solid line is the historical equal phage density line. Thin horizontal black line at the centre divides the system into top and bottom halves. In all three competitions, the historical equal phage density lines (cyan) appear to be flatten over time. Phages *appear* to be neutral at short timescale once *near* steady state conditions are attained at the interface between the respective phage dominated regions. This is a universal rule that is observed in all two-dimensional competitions for all models in this research provided that the competing phages have equal steady state plaque front expansion speeds. Rich dynamics are observed behind the plaque front with reference to the historical equal phage density line (cyan). (a) to (b) In competition D-E, the phage *E* dominated region is expanding into the phage *D* dominated region below the historical equal phage density line. (c) to (d) In competition E-F, the phage *E* dominated region is expanding into the phage *F* dominated region above the historical equal phage density line. (e) to (f) In competition D-F, two phages are almost neutral, with the phage *D* dominated region expanding slightly into the phage *F* dominated region above the historical equal phage density line. Adsorption rates and second phase infection progression rates of competing phages are mentioned in the table F1, and all remaining parameters are mentioned in table 1.

**Figure 9.**
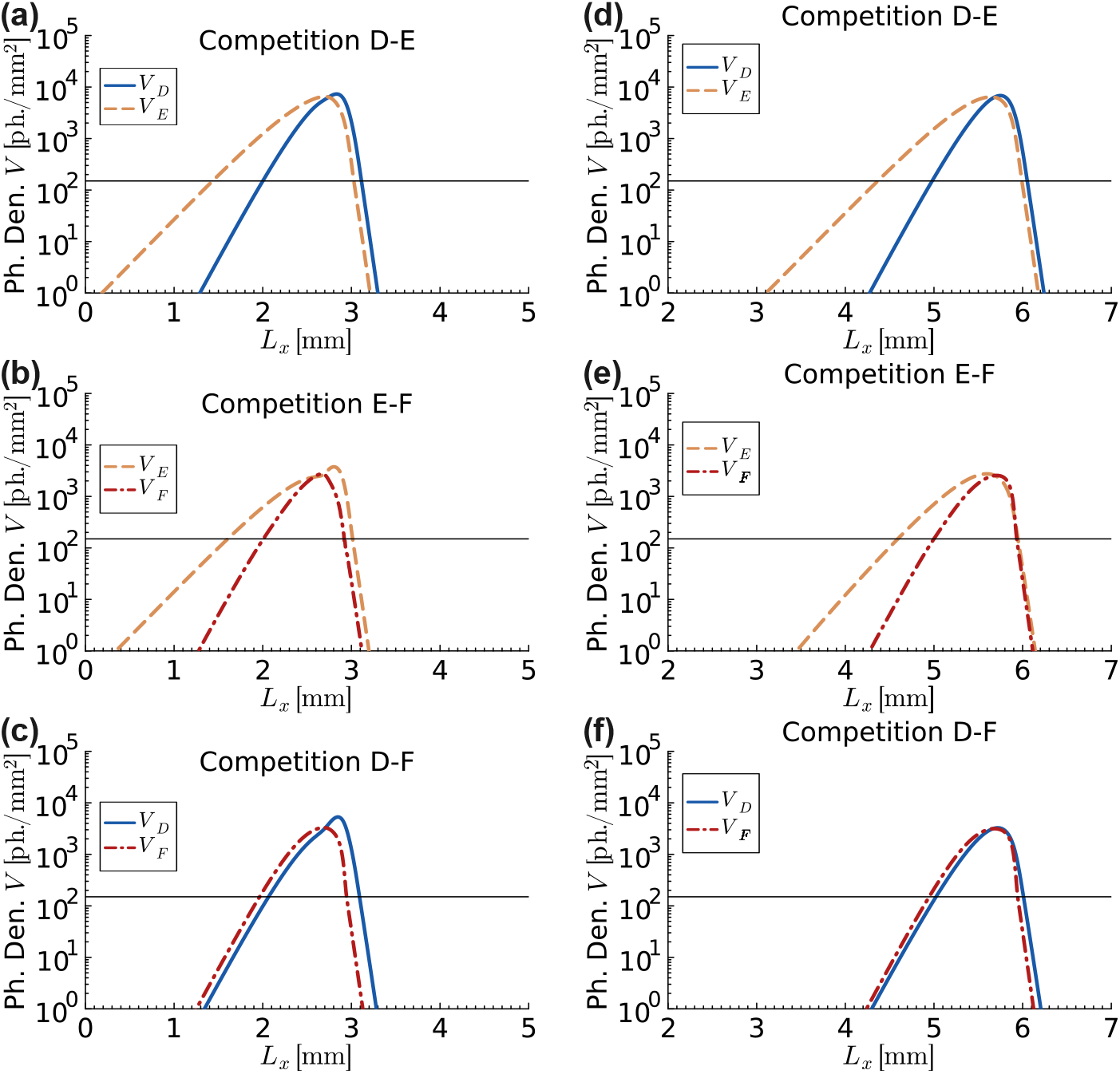
Density profiles of phages competing in a two-dimensional uniform bacterial lawn at a short timescale for the phages with equal steady state plaque front expansion speeds of 5 × 10^−3^ mm/min in isolation for the model with superinfection and discreteness threshold. Phages *D, E* and *F*, having equal steady state plaque front expansion speeds of 5 × 10^−3^ mm/min in isolation, are competed pair-wise in two-dimensional uniform bacterial lawn (fig. 8). Phage density profiles during competition are extracted along the expansion axis (x-coordinate), evaluated at the point where the combined density of the two competing phages is maximal on the equal phage density line (i.e., cyan cross in fig. 8). Snapshots of these profiles are shown at time 50 minutes (left column) and 650 minutes (right column). (a–c) In all competitions, the phage with the lower adsorption rate has a density profile that leads that of the phage with the higher adsorption rate. Consequently, the second phage in each pairing (E in D-E, F in E-F, and F in D-F) initially exhibits a fitness advantage. (d–f) Over time, the density profiles progressively crowd as the lagging phage catches up to the leading one. This convergence of profiles leads to a gradual flattening of the historical equal phage density lines (cyan) in Fig. 8, giving rise to an apparent short-timescale neutrality at the expansion front. Adsorption rates and second phase infection progression rates of competing phages are mentioned in the table F1, and all remaining parameters are mentioned in table 1.

**Figure 10.**
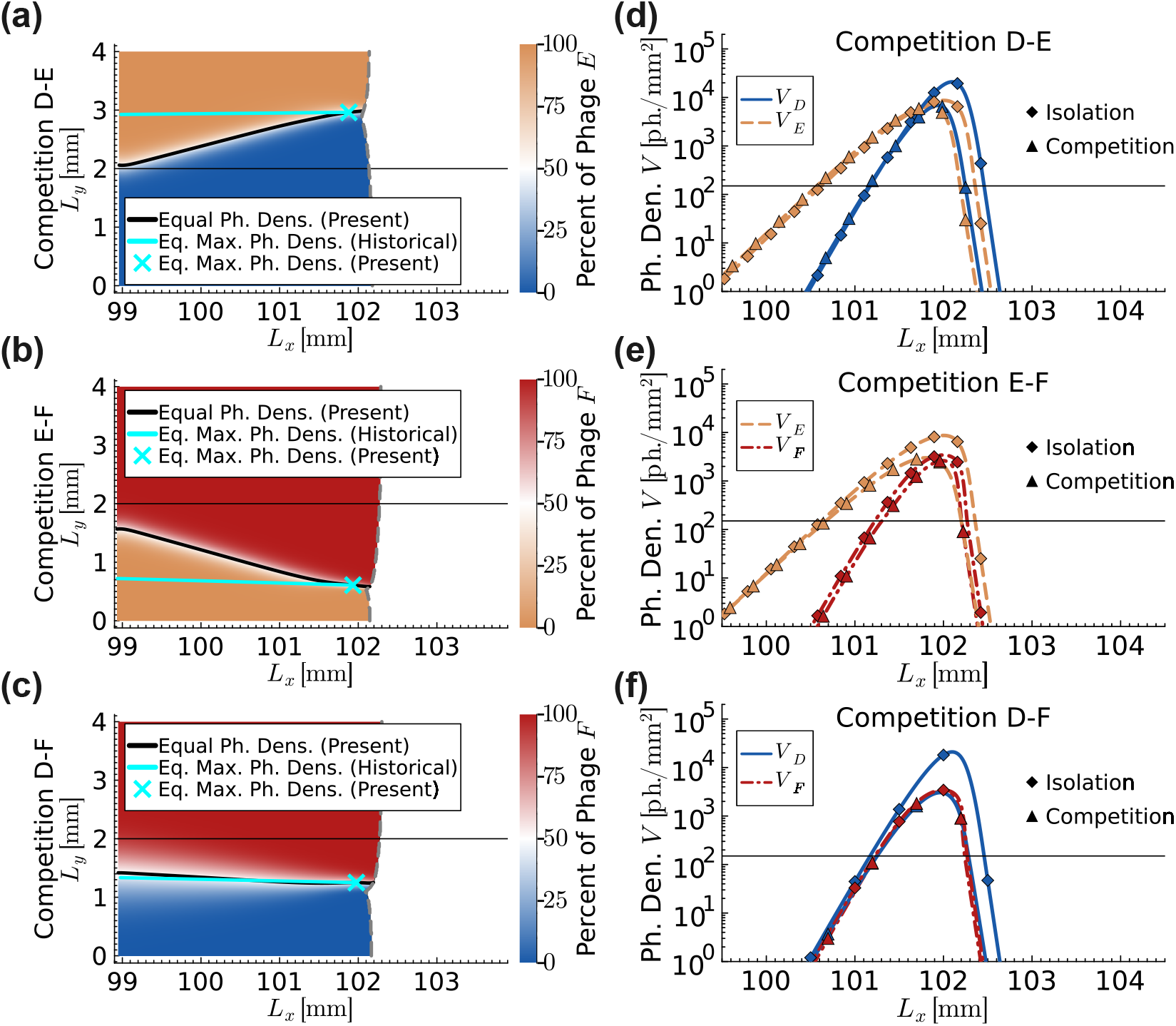
Population dynamics of phages competing in a two-dimensional uniform bacterial lawn at a long timescale for the phages with equal steady state plaque front expansion speeds of 5 × 10^−3^ mm/min in isolation for the model with superinfection and discreteness threshold. Phages *D, E* and *F*, having equal steady state plaque front expansion speeds of 5 × 10^−3^ mm/min in isolation, are competed pair-wise in two-dimensional uniform bacterial lawn. Snapshots of the simulations are given at time 20000 minutes. Different phage dominated regions are shown by their respective colour labels (blue colour for phage *D*, orange colour for phage *E* and red colour for phage *F*). Vertical gray dashed line is the plaque front. Black solid line is the instantaneous equal phage density line and cyan solid line is the historical equal phage density line. Thin horizontal black line at the centre divides the system into top and bottom halves. (a) to (c) In all three competitions, the historical equal phage density lines (cyan) gradually deviated away from the thin horizontal black line over long timescale, indicating that the phages are not neutral over the long timescale. (d) to (f) Curves with diamond markers show phage density profiles in isolation. Curves with triangular markers show phage density profiles in competition along the expansion axis (*x*-coordinate), evaluated at the point where the combined density of the two competing phages is maximal on the equal phage density line (i.e., cyan cross in left column). The thin horizontal black line marks the discreteness threshold for phage density, *θ* = 150 ph.*/*mm. Below this line, phages are effectively absent. In all competitions, the phage that attains maximum density ahead of the other phage eventually wins on the long timescale. Adsorption rates and second phase infection progression rates of competing phages are mentioned in the table F1, and all remaining parameters are mentioned in table 1.

We initially chose for the competition phages D, E and F, as illustrated in fig. 5a. As outlined in the previous section, these phages are characterized by the same plaque front speed in isolation, therefore the front position far away from the interface is expected to be aligned. At the interface between the two populations, we can see that the front recedes (prominently in fig. 10), as a result of the slow-down in expansion due to the shared amount of susceptible cells among the two competing phages and the finite phage discreteness threshold. When a phage strain replicates in isolation, all cells are available to support its replication, allowing its density to quickly exceed the phage discreteness threshold. However, in competition, the resource cells are shared between the two phage strains. This limits the number of cells available to each phage strain to replicate. As a result, each phage population struggles to reach densities above the phage discreteness threshold which slows down the plaque front expansion speed at the interface.

We can also observe over time that while the front away from the interface remains flat, one population takes over the other transversally. This is a different type of invasion compared to what has been previously studied, where one population takes over the other because of different expansion speeds [82]. Later in this paper, we show that when competing phages have different expansion speeds, the faster-expanding phage dominates in side-by-side competitions (fig. 13). Interestingly, we find that the outcome of these competitions in 2D is very different from the outcome in 1D. In the latter, E the most competitive phage (outcompeting both D and F), F was intermediate and D was the least competitive (being outcompeted by both E and F). In 2D, we find that F is the most competitive, followed by D, while E becomes the least competitive, meaning that not only the specific value of phage fitness is affected by the dimensionality of the system, but even the binary outcome of which phage will win in a pairwise competition is.

This behaviour can be understood by examining the relative positions of the phage density profiles across different timescales. Figure 9 shows that near the start of the short-timescale simulations, the density profiles of phages with lower adsorption rates are ahead. This ordering is consistent with the steady-state profiles observed in isolation (fig. 10d–f). Such a transient ordering confers a short-timescale fitness advantage to phages with lower adsorption rates, which manifests as an upward inclination of the historical equal phage density line.

As the competition proceeds, however, the offsets between the competing density profiles gradually decrease, leading to a crowding of the profiles at the expansion front. Consequently, the historical equal phage density line progressively flattens, creating the appearance of neutrality between the competing phages on short timescales.

However, this apparent neutrality does not persist at longer times. As the simulations extend into the long-timescale, the ordering of the phage density profiles can reverse. Figure 10d–f shows that in the competitions E–F and D–F, the initially lagging phage F eventually catches up to and overtakes the initially leading phages (E and D). This reversal in profile ordering confers a long-timescale fitness advantage to phage F, causing the historical equal phage density line to bend downward and ultimately leading to the dominance of phage F in these competitions on long timescale.

Figure 11 illustrates the cross-sections of figure 10 taken along *y*-axis through the location marked by the cyan cross (*L*_*x*_ ≈ 102.0 mm) at time 20000 minutes. This data shows that the outcome of the 2D competition cannot easily be predicted by the phage with the higher density away from the interface (panels b-c, for instance). Indeed, the time-dependent profile of the transversal invasion front (fig. 12) indicates that the outcome of the competition is the result of a non-trivial interplay between phage population dynamics and relaxation to steady-state, making it challenging to pinpoint the exact reason of why one phage outcompetes another.

**Figure 11.**
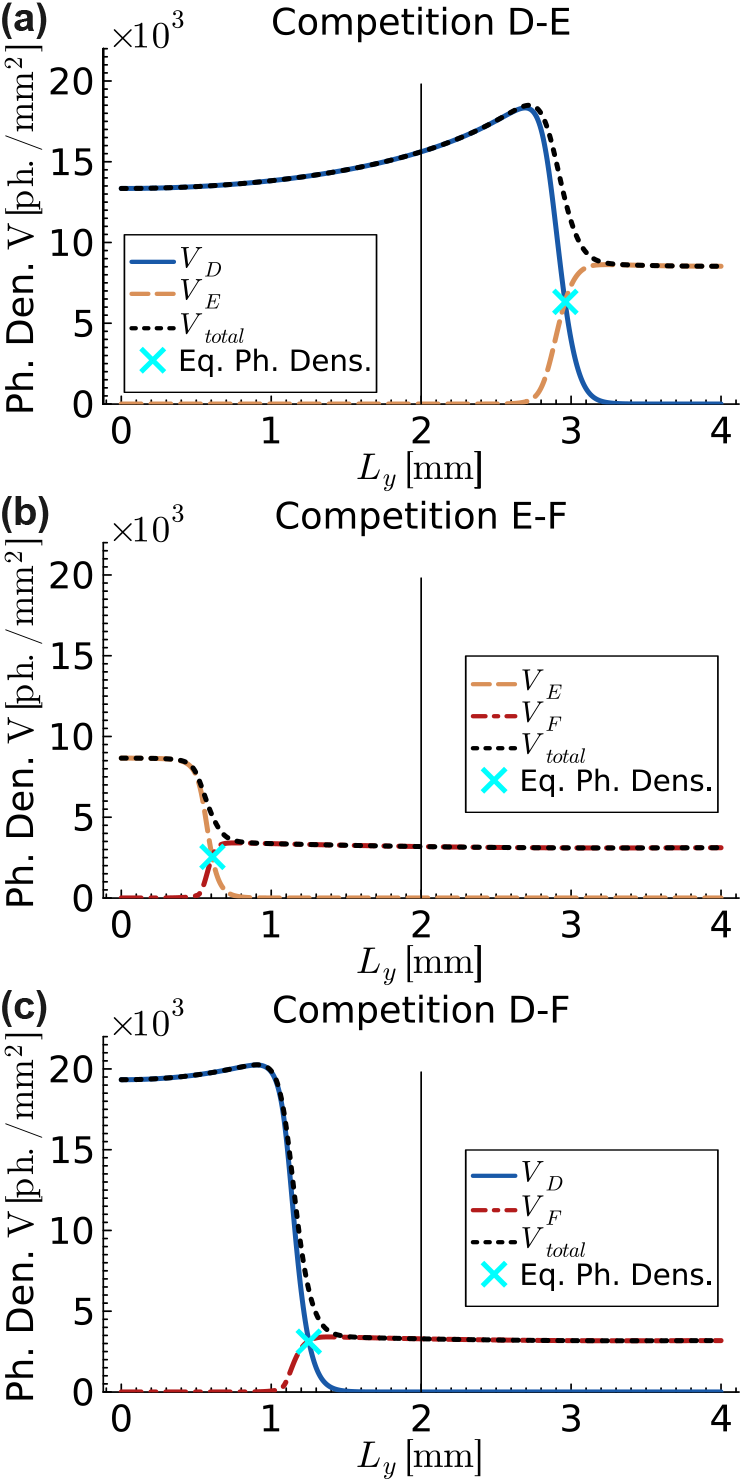
Cross-sectional view of phage density profiles for the phages competing in a two-dimensional uniform bacterial lawn at a long timescale at the points where the combined density of the two competing phages is maximal on the equal phage density line (*i.e*. cyan cross markers in figure 10) for the phages with equal steady state plaque front expansion speeds of 5 × 10^−3^ mm/min in isolation for the model with superinfection and discreteness threshold. Phages *D, E* and *F*, having equal steady state plaque front expansion speeds of 5 × 10^−3^ mm/min in isolation, are competed pair-wise in two-dimensional uniform bacterial lawn. Different phage densities are shown by their respective colour lines (blue solid line for phage *D*, orange dash line for phage *E* and red dotted line for phage *F*). Figure shows phage density profiles along *y*-coordinate at the location around *L*_*x*_ ≈ 102.0 mm at time 20000 minutes as the plaque front passed through the cross section. The thin vertical black solid line at the centre divides the system into left and right halves. (a) and (c) Phage *D* densities peak near the equal phage density points (cyan cross markers), reflecting a slower plaque front progression in competition compared to isolation (see figures 10d to 10f). Adsorption rates and second phase infection progression rates of competing phages are mentioned in the table F1, and all remaining parameters are mentioned in table 1.

**Figure 12.**
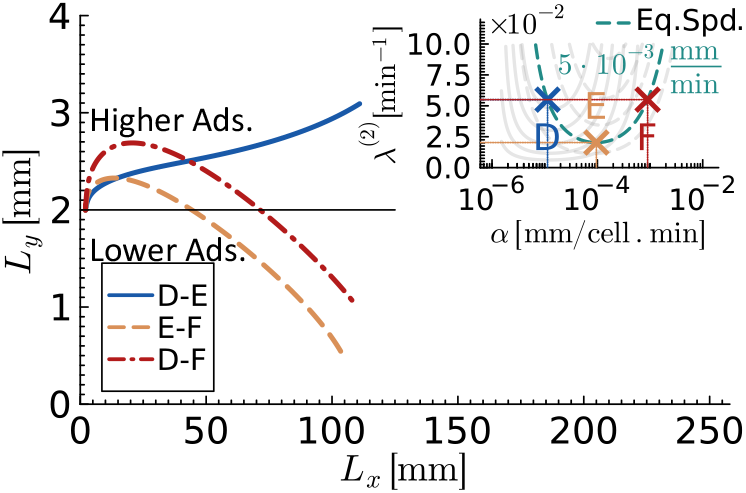
Historical equal maximum phage density lines in a two-dimensional uniform bacterial lawn at a long timescale for the phages with equal steady state plaque front expansion speeds of 5 × 10^−3^ mm/min in isolation for the model with superinfection and discreteness threshold. Simulations are initiated such that the top half of the system contained only the phage with higher adsorption rate, while the bottom half of the system contained only the phage with lower adsorption rate. The plot shows that the phage with lower adsorption rate always win at short timescales, however, over long timescales the trend may shift. In competition D-E, the phage *D* wins on long timescale. In competitions E-F and D-F, phage F wins on long timescale. Adsorption rates and second phase infection progression rates of competing phages are mentioned in the table F1, and all remaining parameters are mentioned in table 1.

Given these results, we expect that neutral competition in 1D is not predictive of neutral competition in 2D. Fig. 13 confirms our expectation by analyzing the competition between phages U, W and F as defined in fig. 6. Phages W and F are neutral in 1D competition, but are characterized by a higher expansion speed for phage F in isolation. As expected, we find that the front associated with phage F bulges out and, over time, takes over the front. Even phages U and F, which are characterized by neutrality in 1D competition and the same expansion speed in isolation, exhibit a transversal invasion speed in favor of phage U at short time (fig. 13) and phage F at long times (fig. 14, fig. 15).

**Figure 13.**
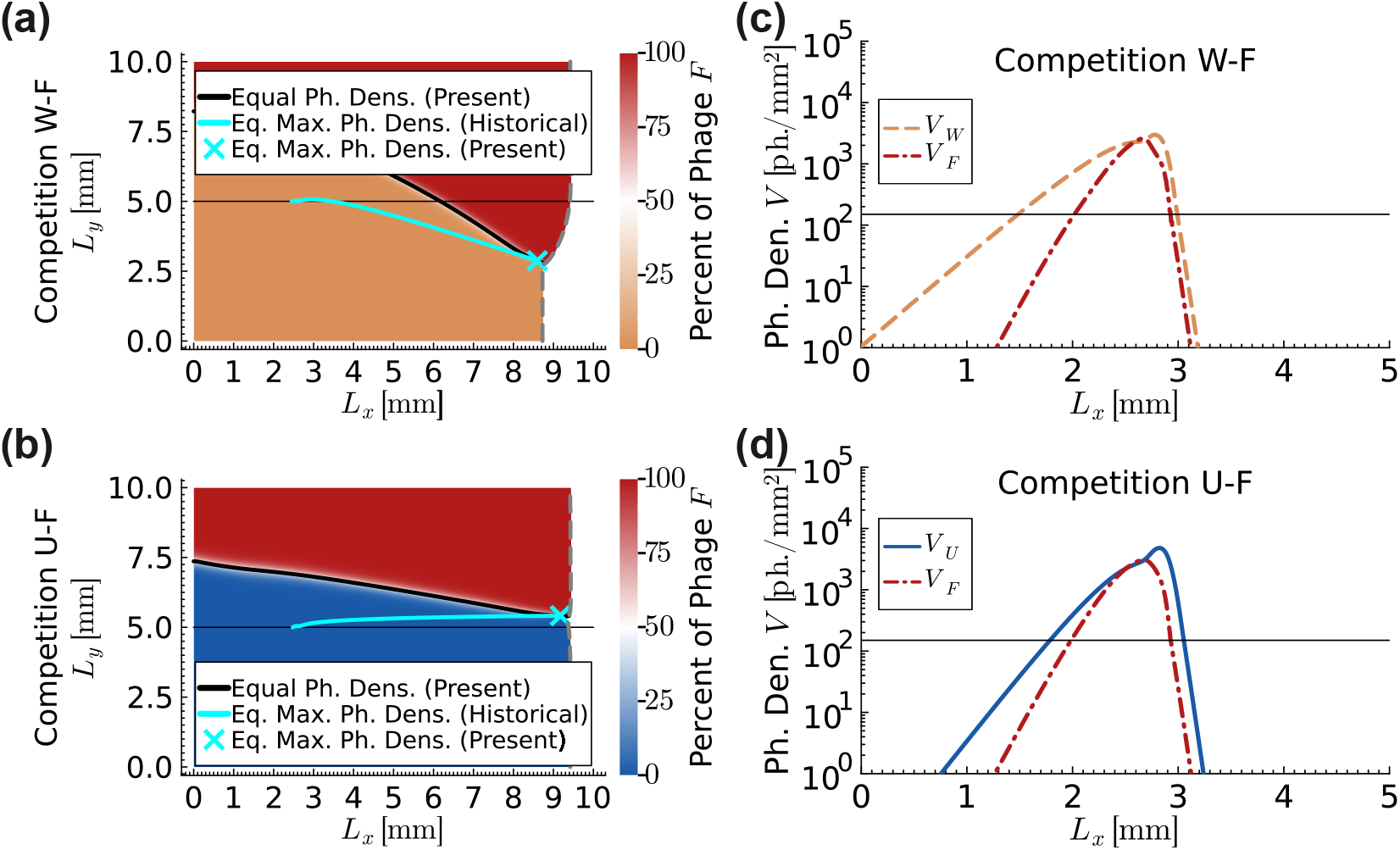
Population dynamics of phages competing in a two-dimensional uniform bacterial lawn at a short timescale for the phages that are neutral when competing in one-dimensional uniform bacterial lawn for the model with superinfection and discreteness threshold. Phages *U, W* and *F*, neutral when compete in one-dimensional uniform bacterial lawn, are competed pair-wise in two-dimensional uniform bacterial lawn. Phages *U* and *F* also have equal steady state plaque front expansion speeds of 5 × 10^−3^ mm/min in isolation, while the steady state plaque front expansion speed of phage *W* is 4.486 × 10^−3^ mm/min in isolation. Snapshots of the simulations are given at times 1350 (left column) and 50 minutes (right column). Different phage dominated regions are shown by their respective colour labels (blue colour for phage *U*, orange colour for phage *W* and red colour for phage *F*). Vertical gray dashed line is the plaque front. Black solid line is the instantaneous equal phage density line and cyan solid line is the historical equal phage density line. Thin horizontal black line at the centre divides the system into top and bottom halves. (a) In competition W-F, the historical equal phage density lines (cyan) continues to move downward because phage *F* has a higher steady state plaque front expansion speed than phage *W*. (b) In competition U-F, the historical equal phage density lines (cyan) appear to be flatten over time. Phages *appear* to be neutral at short timescale once *near* steady state conditions are attained at the interface between the respective phage dominated regions. This is a universal rule that is observed in all two-dimensional competitions for all models in this research provided that the competing phages have equal steady state plaque front expansion speeds. Rich dynamics are observed behind the plaque front with reference to the historical equal phage density line (cyan). (a) and (c) In competition W-F, the phage *W* dominated region is expanding into the phage *F* dominated region above the historical equal phage density line. (b) and (d) In competition U-F, the phage *U* dominated region is expanding into the phage *F* dominated region above the historical equal phage density line. Adsorption rates and second phase infection progression rates of competing phages are mentioned in the table F1, and all remaining parameters are mentioned in table 1.

**Figure 14.**
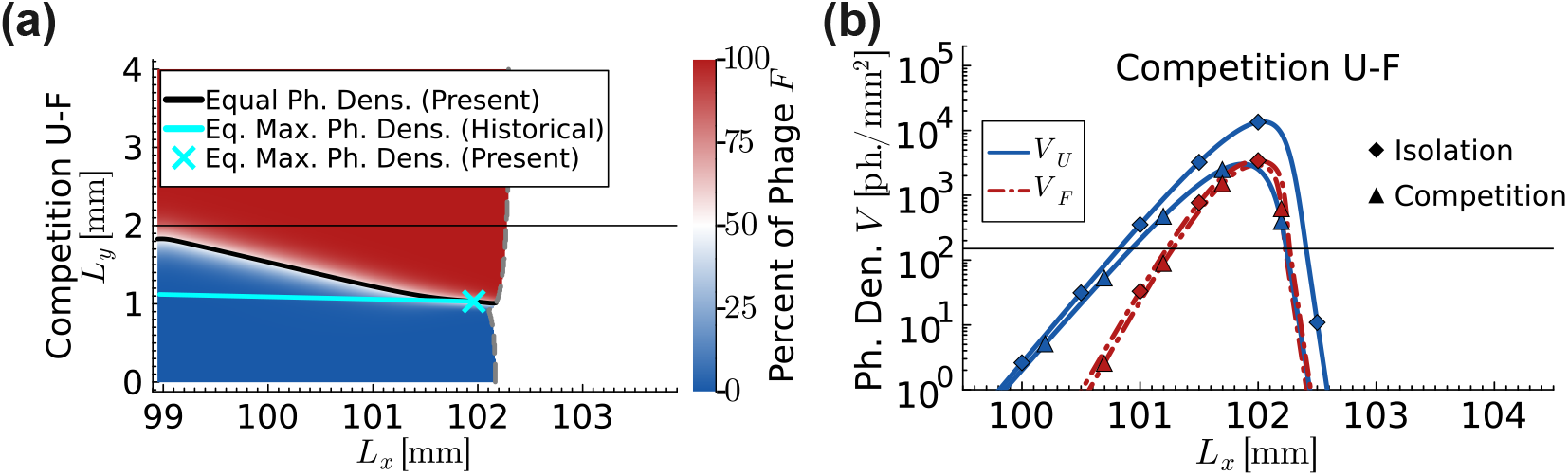
Population dynamics of phages competing in a two-dimensional uniform bacterial lawn at a long timescale for the phages that are neutral when competing in one-dimensional uniform bacterial lawn and have equal steady state plaque front expansion speeds of 5 × 10^−3^ mm/min in isolation for the model with superinfection and discreteness threshold. Phages *U* and *F*, neutral in one-dimensional competitions and have equal steady state plaque front expansion speeds of 5 × 10^−3^ mm/min in isolation, are competed in two-dimensional uniform bacterial lawn. Snapshot of the simulation is given at time 20000 minutes. Different phage dominated regions are shown by their respective colour labels (blue colour for phage *U* and red colour for phage *F*). Vertical gray dashed line is the plaque front. Black solid line is the instantaneous equal phage density line and cyan solid line is the historical equal phage density line. Thin horizontal black line at the centre divides the system into top and bottom halves. (a) In the competition, the historical equal phage density line (cyan) gradually deviated away from the thin horizontal black line over long timescale, indicating that the phages are not neutral over the long timescale. (b) Curves with diamond markers show phage density profiles in isolation. Curves with triangular markers show phage density profiles in competition along the expansion axis (*x*-coordinate), evaluated at the point where the combined density of the two competing phages is maximal on the equal phage density line (i.e., cyan cross in left column). The thin horizontal black line marks the discreteness threshold for phage density, *θ* = 150 ph.*/*mm. Below this line, phages are effectively absent. In all competitions, the phage that attains maximum density ahead of the other phage eventually wins on the long timescale. Adsorption rates and second phase infection progression rates of competing phages are mentioned in the table F1, and all remaining parameters are mentioned in table 1.

**Figure 15.**
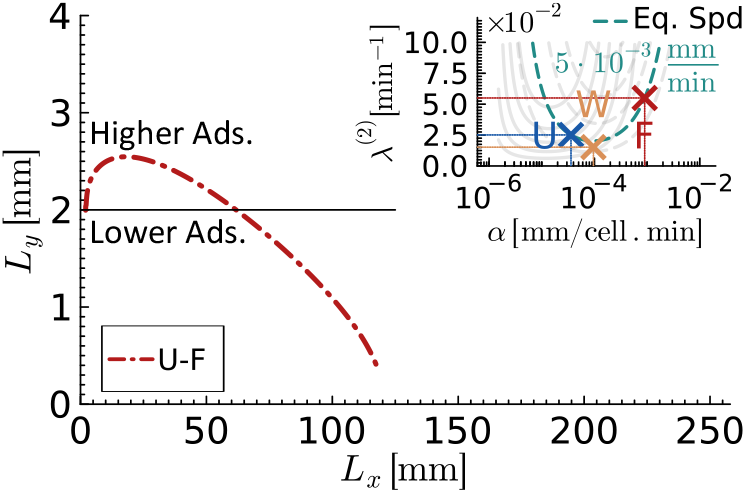
Historical equal maximum phage density lines in a two-dimensional uniform bacterial lawn at a long timescale for the phages that are neutral when competing in one-dimensional uniform bacterial lawn and have equal steady state plaque front expansion speeds of 5 × 10^−3^ mm/min in isolation for the model with superinfection and discreteness threshold. Simulation is initiated such that the top half of the system contained only the phage with higher adsorption rate, while the bottom half of the system contained only the phage with lower adsorption rate. The plot shows that the phage with lower adsorption rate wins at short timescales, however, over long timescales the trend shifts and the phage *F* wins on long timescale. Adsorption rates and second phase infection progression rates of competing phages are mentioned in the table F1, and all remaining parameters are mentioned in table 1.

## 4. DISCUSSION

Evolutionary fitness is typically defined as an individual’s survival and reproductive success in the environment it inhabits [83]. Because the quantity is by definition context-dependent, its mathematical expression and experimental quantification have traditionally been challenging, often requiring assumptions that can easily break. While this limitation of fitness is recognized and even exploited when investigating traits selected for by different environments [84–88], it is not fully appreciated when comparing situations that, superficially, appear equivalent, leading to potentially inappropriate comparisons between models and experiments.

Phage and other viral populations are a particularly interesting system in which to explore this question, since the growth rate is not an independent quantity, but emerges from the infection dynamic as a function of the life history parameters of the virus, e.g., adsorption rate, lysis time and burst size [34, 35, 38]. We have previously shown that this coupling leads to unexpected differences between phage effective growth rate in isolation and in direct competition even in well-mixed turbidostat scenarios, where the available resources (host cells) are held constant [24].

Here, we investigate this question in depth in the context of spatial range expansions, where both growth and dispersal contribute to the fitness of a population, and the emerging *expansion speed* has been typically used as the appropriate measure for fitness, and shown, at least in microbial colonies, to be an excellent predictor for competition outcome [82].

By taking advantage of the internal infection dynamic that generates a trade-off between adsorption rate and lysis rate, we identify the combinations in parameter space that result in the same expansion speed in isogenic phage populations. This allows us to systematically assess whether two phage populations with different infection parameters but the same expansion speed in isolation behave neutrally in a direct competition.

To analyze the phage population dynamics, we have developed a set of spatiotemporal models based on coupled systems of non-linear partial differential reaction-diffusion equations. The models were designed to simulate replication of one or two lytic phage strains in a microbial lawn consisting of one bacterial strain. We have assumed the presence of a post-adsorption superinfection exclusion mechanism, which is common across many phages, and accommodated phage discreteness. Variability in lysis time was captured by implementing two subsequent phases of infection, each with exponentially distributed rates, leading to a hypoexponential distribution of total lysis time.

The model is sufficiently flexible to be easily extended to accommodate: (i) different modes of superinfection or superinfection exclusion (partly explored in Appendix C) [24, 89], (ii) distinct lysis time distributions by decreasing or increasing the number of intermediate infection phases and changing their rates [53], (iii) a time-dependent burst size by introducing a lysis rate and corresponding burst size for each infection phase [37, 90]. While each of these extensions inevitably makes the computation more demanding, the model comes with the advantage of enabling quantification and comparison of all these factors in one single mathematical framework.

Our results show that expansion speed is a good predictor for phage competition outcome on bacterial lawns only if the discreteness of the phage population is neglected. In this case, the phage fields are allowed to reach unrealistically low values at the tip of the expansion, which is the part of the wave that drives the expansion forward and sets the velocity [61]. Because the value of phage density is so small in this area, competition for resources plays no role, leading to identical results in competition and isolation. If we take into account phage discreteness, then phage replication occurs deeper in the expanding wave, where more phages are present and resource competition becomes relevant, manifesting as a reduction in the expansion speed compared to the same phage grown in isolation (fig. 2 vs. fig. D1).

The significant role of resource competition leads to interesting and non-trivial results when two distinct phages compete for the same space, as the ensemble of phages expected to be neutral from speed in isolation is different from the ensemble of phages that turns out to actually be neutral in direct competition (fig. 6). This observation implies not only that the speed in isolation cannot predict competition outcome, but also that the expansion speed of one phage is altered by the presence of another phage in a non-trivial way. Using phage F as reference, Fig. 6 shows that the equal-speed curve intersects the 1D isofitness curve in two points, so that the isofitness curve is lower in the middle and higher outside. This implies a strong context-dependent fitness effect so that phages with intermediate adsorption rates (between U and F) will be slowed down by phage F and, vice versa, phages with very low or very high adsorption rates (below U or above F) will slow down phage F.

The observation of this context-dependent effect in which the expansion speed of a phage depends on the presence of competing phages raises questions around the transferability of the isofitness curves. We find that while all the points laying on the black line in fig. 6 are neutral to phage F in direct competition, they are not necessarily neutral with respect to each other (for instance U and W in fig. 7). This implies that fitness cannot be measured in reference to a single competing phage, but strongly depends on the nature of the *two* (or more) competing phages, so that, in principle, for each phage strain we could draw its own isofitness line. For instance, if we were to draw the isofitness line for phage U, it would cross the isofitness line for phage F at F (U and F are neutral with respect to each other) but would sit above the isofitness lines for F in between U and F (as we know that U outcompetes W). The lack of transitive property of competing fitness leads to unexpected rock-scissor-paper dynamics, where U would outcompete phage X (located as in the fig 16), which would outcompete phage F, which would outcompete phage Y (as in fig 16b), which would outcompete phage U, for instance. We call this type of selection *ecology-dependent*, since the viral trait selected for would depend on the phenotypic make-up of the whole population. In terms of evolutionary experiments, this would lead to evolutionary trajectories that heavily depend on the relative frequencies of concurrent mutant clones, raising interesting questions regarding the robustness and predictability of evolutionary scenarios.

**Figure 16.**
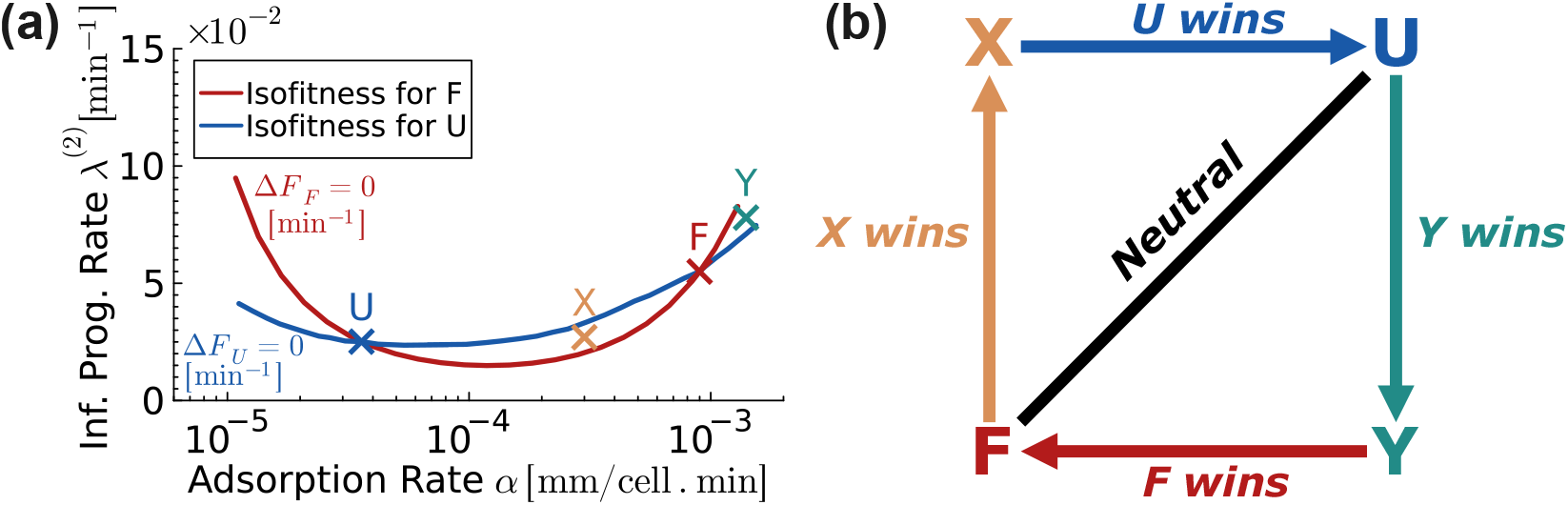
Rock-scissor-paper dynamics and schematic of isofitness curves for the one-dimensional two phage strains model for phages *U* and *F*. (a) For two phage strains models, the two isofitness curves represent sets of phage life-history parameters for which the relative fitness is zero with respect to phage *U* (Δ*F*_*U*_ = 0 min^−1^, blue solid line) and phage *F* (Δ*F*_*F*_ = 0 min^−1^, red solid line), as defined in equation 7. The two intersection points of these curves correspond to phages *U* and *F* which are neutral in one-dimensional competition. As shown, the isofitness curve for phage *U* lies above that of phage *F*. Arbitrary phages *X* and *Y* are located between the two isofitness curves. Schematic illustrate the rock–scissors-paper-like dynamics among phage strains. Phage *U* outcompetes *X*, which outcompetes *F*, which outcompetes *Y*, which outcompetes *U*. Adsorption rates and second phase infection progression rates of phages *U* and *F* are mentioned in the table F1, and all remaining parameters are mentioned in table 1.

Because the spatial population dynamic depends on the type and abundance of the competing phages, the generalizability of 1D results into 2D environments where two phage populations compete side by side is not straightforward: by construction, in this configuration different transversal positions along the expanding front will exhibit different proportions of competing phages (see, for instance, cross-sections in fig. 11), each resulting in a different expansion velocity. This is readily observable in fig. 10, where the front is flat and aligned far away from the interface, as the two competing phages are chosen to have the same expansion speed in isolation, but dips closer to the interface. Interestingly, beyond the effect on the expansion speeds and front profile, the density-dependent nature of competition leads to counter-intuitive observation, where, for instance, the density of one phage is boosted by the nearby presence of a competing phage (phage D in fig. 11a,c). This phenomenon, which emerges in systems with dimensions higher than 1 due to the local slowdown of the plaque front, might contribute to the unexpected result that increasing dimensions (from 1D to 2D) may flip the outcome of a competition (as for instance, phages D and E in fig. 5 vs. fig. 10).

While it is difficult to pin-point simple rules that comprehensively explain the outcome of a given phage competition, we can identify the key element that gives rise to the competitor-dependent selection we observe: the interplay between consumption of shared available resources and phage replication. Since in our model the two competing phage populations do not interact in any direct way, our observations must stem from the indirect interactions that arise from competition for a common resource (bacteria). Importantly, the dynamic emerging from this competition strongly depends on the types of competing phages and not just on their numbers, which is why a phage competing with itself behaves differently from the same phage competing with another phage. In our work, this dynamic is modulated by the trade-off between the phage’s adsorption rate and the lysis time, which measure how efficiently and quickly the population converts resources, respectively. Going beyond phages, we might expect to see similar behaviour in populations that compete for the same resource, e.g., same nutrient, but consume it at different rates in exchange for faster or slower replication rates. While previous work has investigated some of the aspects of resource competition in range expansions [91–94], as far as we know the effects of the trade-off between resource consumption and replication rate has not been explored. We speculate that even simple models that accommodate such trade-off might lead to ecology-dependent fitness landscapes similar to what we see here for phages.

## Acknowledgments

We thank Jacopo Marchi and the rest of the Fusco group for useful discussions. This research was financially supported by a Cavendish Scholarship (Sackler Trust Fund) and an ERC Starting Grant/UKRI Horizon Europe Guarantee (EP/Y030141/1, DF). Simulations in this work were performed using resources provided by the Cambridge Service for Data Driven Discovery (CSD3) operated by the University of Cambridge Research Computing Service (www.csd3.cam.ac.uk), provided by Dell EMC and Intel using Tier-2 funding from the Engineering and Physical Sciences Research Council (capital grant EP/T022159/1), and DiRAC funding from the Science and Technology Facilities Council (www.dirac.ac.uk).

## Data Availability

Codes are available at: https://github.com/FuscoLab/2025_phage_fitness_in_spatial_expansions.

