## Supplementary Material for "Resource availability and dimensionality result in ecology-dependent selection in bacteriophage spatial expansions"

### Supplementary information for "Resource availability and dimensionality result in ecology-dependent selection in bacteriophage spatial expansions"

**Hassan Alam and Diana Fusco**

Cavendish Laboratory, University of Cambridge, Cambridge CB3 0HE, United Kingdom

September 2025

#### Appendix A. Supplementary Materials and Methods

*Appendix A.1. Equations of the two phage and one bacterial strain model*

The model is based on the coupled system of the non-linear differential equations that describe the viral and host population dynamics of microbial system. The equations of the spatiotemporal model for the two phage and one bacterial strain model are given below.

$$\frac{\partial B}{\partial t} = -\alpha_1 V_1 \cdot T(V_1) \cdot B - \alpha_2 V_2 \cdot T(V_2) \cdot B \tag{A.1}$$

$$\frac{\partial I_1^{(1)}}{\partial t} = \alpha_1 V_1 \cdot T(V_1) \cdot B - \lambda_1^{(1)} I_1^{(1)} \tag{A.2}$$

$$\frac{\partial I_2^{(1)}}{\partial t} = \alpha_2 V_2 \cdot T(V_2) \cdot B - \lambda_2^{(1)} I_2^{(1)} \tag{A.3}$$

$$\frac{\partial I_1^{(2)}}{\partial t} = \lambda_1^{(1)} I_1^{(1)} - \lambda_1^{(2)} I_1^{(2)} \tag{A.4}$$

$$\frac{\partial I_2^{(2)}}{\partial t} = \lambda_2^{(1)} I_2^{(1)} - \lambda_2^{(2)} I_2^{(2)} \tag{A.5}$$

$$\begin{aligned}\frac{\partial V_1}{\partial t} = {} & \beta_1 \lambda_1^{(2)} I_1^{(2)} - \delta_1 V_1 - \alpha_1 V_1 \cdot T(V_1) \cdot B - \alpha_1 V_1 \cdot T(V_1) \cdot I_1^{(1)} - \alpha_1 V_1 \cdot T(V_1) \cdot I_2^{(1)} \\ & - \alpha_1 V_1 \cdot T(V_1) \cdot I_1^{(2)} - \alpha_1 V_1 \cdot T(V_1) \cdot I_2^{(2)} + D\nabla^2 V_1\end{aligned} \tag{A.6}$$

$$\begin{aligned}\frac{\partial V_2}{\partial t} = {} & \beta_2 \lambda_2^{(2)} I_2^{(2)} - \delta_2 V_2 - \alpha_2 V_2 \cdot T(V_2) \cdot B - \alpha_2 V_2 \cdot T(V_2) \cdot I_1^{(1)} - \alpha_2 V_2 \cdot T(V_2) \cdot I_2^{(1)} \\ & - \alpha_2 V_2 \cdot T(V_2) \cdot I_1^{(2)} - \alpha_2 V_2 \cdot T(V_2) \cdot I_2^{(2)} + D\nabla^2 V_2\end{aligned} \tag{A.7}$$

where, $D$ is the diffusion coefficient and $\nabla^2 = \partial^2/\partial x^2$ for one-dimensional space and $\nabla^2 = \partial^2/\partial x^2 + \partial^2/\partial y^2$ for two-dimensional space.

The model without a phage discreteness threshold is obtained by setting $T(V_i) = 1$ in all equations, where $i = 1, 2$. The model without phage superinfection is obtained by removing all $\alpha_i V_i \cdot T(V_i) \cdot I_j^{(k)}$ terms, where $i, j = 1, 2$.

*Appendix A.2. Numerical Approximations and Boundary Conditions*

The system of partial differential equations is converted to the system of ordinary differential equations by approximating the diffusion term using finite difference method. Space is divided into lattice sites of finite size $\Delta x$ (and $\Delta y$). Each partial differential equation splits into large number of coupled ordinary differential equations. To calculate the spatial derivatives for the diffusion terms, second-order central finite difference formula is used.

$$\left.\frac{d^2 V}{dx^2}\right|_{x_i} \approx \frac{V(x_{i+1}) - 2V(x_i) + V(x_{i-1})}{(\Delta x)^2} \tag{A.8}$$

However, to handle the left boundary, second-order forward finite difference was used,

$$\left.\frac{d^2V}{dx^2}\right|_{x_1} \approx \frac{V(x_3) - 2V(x_2) + V(x_1)}{(\Delta x)^2} \tag{A.9}$$

And, to handle the right boundary, second-order backward finite difference was used.

$$\left.\frac{d^2V}{dx^2}\right|_{x_{\text{end}}} \approx \frac{V(x_{\text{end}}) - 2V(x_{\text{end}-1}) + V(x_{\text{end}-2})}{(\Delta x)^2} \tag{A.10}$$

This allows the phages to smoothly flow out of the system through its boundaries. The same method is used to handle the top and bottom boundaries (derivatives in $y$-direction) for the two-dimensional models.

The resulting set of large number of ordinary differential equations is solved numerically. To keep the solution stable, the maximum time step is limited to $dt_{\text{max}} = \Delta x^2/2D$. A minimum time step of $dt_{\text{min}} = 10^{-300}$ is used to avoid truncation and rounding errors caused by the finite precision of computer arithmetic. Values of densities are also constrained to non-negative values. Note that above conditions are relaxed in some extreme cases when iteration takes unrealistically long time to complete.

Bogacki-Shampine 3/2 (BS3) algorithm is used to solve the equations of the models with phage discreteness threshold as the algorithm is fast but less accurate. However, Dormand-Prince's 5/4 Runge-Kutta (DP5) algorithm is used to solve the equations of the model without phage discreteness threshold as the algorithm is slow but more accurate.

During simulations, after initial transient periods, both the maximum phage density at the plaque front and the plaque front expansion speed approach steady state values. In models with the phage discreteness threshold, these quantities reach constant steady state values after some finite time (see Figure A1). In contrast, in models without a discreteness threshold, they approach steady state limits only asymptotically. The method used to estimate these steady state limits is described in Section Appendix A.3. To measure the size (radius) and speed of the plaque, we define the plaque front as the location of the maximum density of the total infected cells.

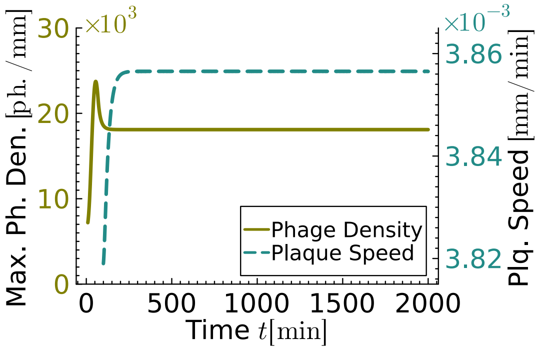


Figure A1: **Maximum phage density at the plaque front and plaque front expansion speed approach steady state values after a finite time for the models with phage discreteness threshold**. The figure shows that with time, maximum phage density at the plaque front and plaque front expansion speed approach steady state values after a finite time for the models with phage discreteness threshold. In the study, plaque front is defined as the location at which the cumulative density of infected cells is maximum. The figure is obtained by using the one-dimensional one phage strain model with superinfection and discreteness threshold. Adsorption rate is $5 \times 10^{-6}$ mm/cell · min, and all remaining parameters are mentioned in table 1.

*Appendix A.3. Method to calculate asymptotic limits for the models without discreteness threshold*

For the models without a phage discreteness threshold, the maximum phage density at the plaque front and the plaque front expansion speed asymptotically approach to steady state values over time. As infinite-time simulations are impractical, these asymptotic limits are determined using the following mathematical approach.

Assume we have a variable $y(x)$ that depends on $x$ and it is approaching its steady state value $y_{ss}$ with $x$. If the asymptotic assumption is true, $y_{ss} - y$ should follow a power-law with $x$.

$$y_{ss} - y \propto x^m$$

$$y_{ss} - y = e^k x^m \tag{A.11}$$

$$\tag{A.12}$$

where $e^k$ is a constant. Let $z \equiv y_{ss} - y$

$$z = e^k x^m \tag{A.13}$$

Take logarithm on both sides,

$$\ln z = \ln (y_{ss} - y) = k + m \ln x \tag{A.14}$$

This implies that the log-log graph between the $(y_{ss} - y)$ versus $x$ is a straight line if $y$ is asymptotically approaching the steady state value $y_{ss}$.

To iterate the value of $y_{ss}$, we guess a value of $y_{guess}$ of the steady state $y_{ss}$, then we plot log-log graph between $(y_{guess} - y_{data})$ versus $x_{data}$, where $x_{data}$ and $y_{data}$ are the numerical data obtain from the simulations. If the guess value $y_{guess}$ is a true steady state value $y_{ss}$, the log-log graph results in a straight line. To numerically calculate the straightness of the graph against a guessed value $y_{guess}$, we fit the straight line equation $\ln (y_{guess} - y) = k + m \ln x$ on the numerical data plot, $\ln (y_{guess} - y_{data})$ versus $\ln x_{data}$, by using regression. Then, we find the sum square error $SSE$ between the fitted straight line and numerical data as follows,

$$SSE = \Sigma |z - z_{data}|^2 \tag{A.15}$$

where, $z = e^k x_{data}^m$ and $z_{data} = y_{guess} - y_{data}$. Therefore, the value of $y_{guess}$ that minimizes the $SSE$ should be the true value of $y_{ss}$. If the resulting log-log graph between $(y_{ss} - y_{data})$ versus $x_{data}$ is a straight line, then the assumption that the variable is asymptotically approaching a steady state value is true.

*Appendix A.4. Initial configurations for one-dimensional simulations*

One-dimensional simulations start with the initial condition that uninfected bacterial cells are uniformly distributed throughout the system with a constant density of 1000 cells/mm, and initial densities of infected bacterial cells are taken to be zero.

Initially, phage densities follow very narrow normal distributions, resembling delta functions, with mean $\mu = 0$ ph./mm and standard deviation $\sigma = 0.01$ ph./mm representing phage particle(s) of each strain initially positioned at the left boundary of the systems. For the models with the phage discreteness threshold, 150 phage particles of each strain are distributed according to the normal distribution. In contrast, models without the phage discreteness threshold initialise with only one phage particle per strain.

The length of the system is $L_x = 10$ mm for short timescale simulations, and the length of the system is $L_x = 5$ mm for long timescale simulations.

*Appendix A.5. Initial configurations for two-dimensional simulations*

Two-dimensional simulations are initialised with two side-by-side plaques having aligned steady-state fronts, expanding from left to right across the systems.

In the simulations, we want to examine the relative fitness of phages having equal steady state plaque front expansion speeds. Therefore, we need the system to be in the steady state right from the start of these simulations. The two-dimensional initial condition data is obtained by replicating the one-dimensional steady state data in $y$-direction.

For the short timescale simulations, one-dimensional steady state data of the phage with lower adsorption rate is replicated in range 0 mm $< L_y <$ 5 mm (*i.e.*, bottom half of the two-dimensional system) and one-dimensional steady state data of the phage with higher adsorption rate is replication in range 5 mm $< L_y <$ 10 mm (*i.e.*, top half of the two-dimensional system) while the length of the system in $x$ direction is $L_x = 10$ mm. For the long timescale model, the corresponding ranges are 0 mm $< L_y <$ 2 mm and 2 mm $< L_y <$ 4 mm, respectively, while $L_x = 5$ mm.

The steady state one-dimensional data is also shifted to align the peaks of the density of total infected cells (*i.e.*, plaque fronts). The distribution of phages behind the plaque front also effects the plaque front expansion speed. Therefore, the plaque fronts are initially located at the position $x = 2\frac{2}{3}$ mm, to retain sufficient steady state data behind the plaque front to account for the effects of phage diffusion. Since the plaque fronts are shifted toward the left boundary, extrapolation is required at the right boundary to avoid introducing fictitious zero or undefined values. The phage and infected cell densities are assumed to decay exponentially near the right boundary. Accordingly, the last 40 data points of each variable are used to fit the function $y = e^{-ax+b}$, which is then employed to extrapolate the missing values at the right boundary after shifting the plaque front.

For models with phage discreteness threshold, for long timescale competitions, the two-dimensional plaques are expanded to a size of 100 mm which takes about four orders of magnitude of simulation time $O(10^4)$ in minutes to attain this size. It is critical to obtain very accurate values of plaque front expansion speeds for long timescale competitions; otherwise, only a minor difference between the plaque front expansion speeds of competing phages would accumulate over time to give observable lead to the phage with slightly higher speed in two-dimensional plaques. Therefore, after selecting a set of phages (that is, phages $D$, $E$, $F$) against a fixed plaque front expansion speed in isolation (*i.e.*, $5 \times 10^{-3}$ mm/min), their one-dimensional simulations in isolation are performed again till the plaque size of 1000 mm to ensure that steady state plaque front expansion speeds are accurately achieved. Values of adsorption rates of the phages $D$ and $F$ are refined by using simple linear interpolation technique until the speeds of all three phages in a set differ only by the order of $O(10^{-15})$ mm/min.

For models without a phage discreteness threshold, performing long timescale simulations is not straightforward, as numerical computations are limited by machine precision when handling extremely small values. Therefore, in this paper, only short timescale simulations are performed for such models. To obtain *near* steady state data for such models, one-dimensional single phage simulations were performed for phages having *near* equal plaque front expansion speeds (*i.e.*, phages $D$, $E$ and $F$) in which plaques are allowed to evolve for 1200 minutes. By the end of all simulations, the plaque front expansion speeds were above 99.5% of their asymptotic limits, ensuring that the final states of the simulations were reasonably close to their steady state limits. In these simulations, the length of the systems was taken to be 20 mm to ensure continuity in numerical solutions because the position of the plaque front in the steady state data was shifted towards the left boundary for the two-dimensional simulations. This adjustment was necessary to avoid fictitious zero-valued or undefined elements at the right boundary, ensuring the grid size of the steady state data match the desired grid size in $x$-direction of the two-dimensional simulations.

##### *Appendix A.6. Repeated reinitialization procedure by shifting plaque for long timescale simulations*

Simulation domain sizes need to be sufficiently small to minimise the computational time. A discrete treadmill scheme is used to obtain steady state data when using long timescale simulations for one-dimensional and two-dimensional models with phage discreteness threshold. In this scheme, as the plaque front reaches near the right boundary of the system, the complete system is shifted leftwards near the left boundary of the system. Since the plaque fronts are shifted toward the left boundary, extrapolation is required at the right boundary to avoid introducing fictitious zeros or undefined values. The phage and infected cell densities are assumed to decay exponentially near the right boundary. Accordingly, the last 40 data points of each variable are used to fit the function $y = e^{-ax+b}$, which is then employed to extrapolate the missing values at the right boundary after shifting the plaque front. Table A1 gives the treadmill durations used during the transient and *near* steady-state phases of the long timescale simulations.

Table A1: **Steady state and transient run periods used in treadmill-based long timescale simulations.**

| **Plaque Front Speed [mm/min]** | **Non Steady State Period [min]** [(a)] | ***Near* Steady State Period [min]** [(b)] |
|---|---|---|
| $5.0 \times 10^{-3}$ | 500 | 200 |
| $4.4862 \times 10^{-3}$ | 550 | 225 |
| $6.0 \times 10^{-3}$ | 400 | 150 |
| $7.0 \times 10^{-3}$ | 350 | 150 |
| $8.0 \times 10^{-3}$ | 300 | 100 |
| $4.0 \times 10^{-3}$ | 625 | 250 |
| $5.6412 \times 10^{-3}$ | 450 | 150 |
| $9.2572 \times 10^{-3}$ | 300 | 100 |

[(a)] **One-dimensional simulations often start with non steady state initial conditions**. This duration is used in the first run of treadmill simulations when steady state initial conditions are unknown because plaque front expansion speed is slow during initial transient period.
[(b)] This duration is used for treadmill-based long timescale simulations in *near* steady state.

#### Appendix B. Additional Competitions For The Model With Superinfection And Discreteness Threshold At Different Steady State Plaque Front Expansion Speeds

To further explore the parameter space, additional competitions for the model with superinfection and discreteness threshold have been performed. Phages $D$, $E$, and $F$ are selected from the different isofitness curves (equal steady state plaque front expansion speeds) at speeds $8 \times 10^{-3}$, and $4 \times 10^{-3}$ mm/min. Results from the competitions are shown in this section.

Results from these additional competitions are mostly in agreement with the results discussed in the main text with a few exceptions that are discussed below.

In one-dimensional uniform bacterial lawn competition for the two phage strains model with superinfection and discreteness threshold, phages $E$ win in competition E-F when the steady state plaque front expansion speed was $5 \times 10^{-3}$ mm/min (see fig. 5). However, when steady state plaque front expansion speed is increased to $8 \times 10^{-3}$ mm/min, phage F wins in the competition E-F (see figure B1).

For the competition D-E of phages having steady state plaque front expansion speed of $4 \times 10^{-3}$ mm/min, figure B9(b) shows that the phage $E$ marginally wins in the two-dimensional competition D-E on contrary to the other competitions D-E in which phage $D$ consistently won. This might be because here the phage $E$ is not located at the minimum of the isofitness curve.

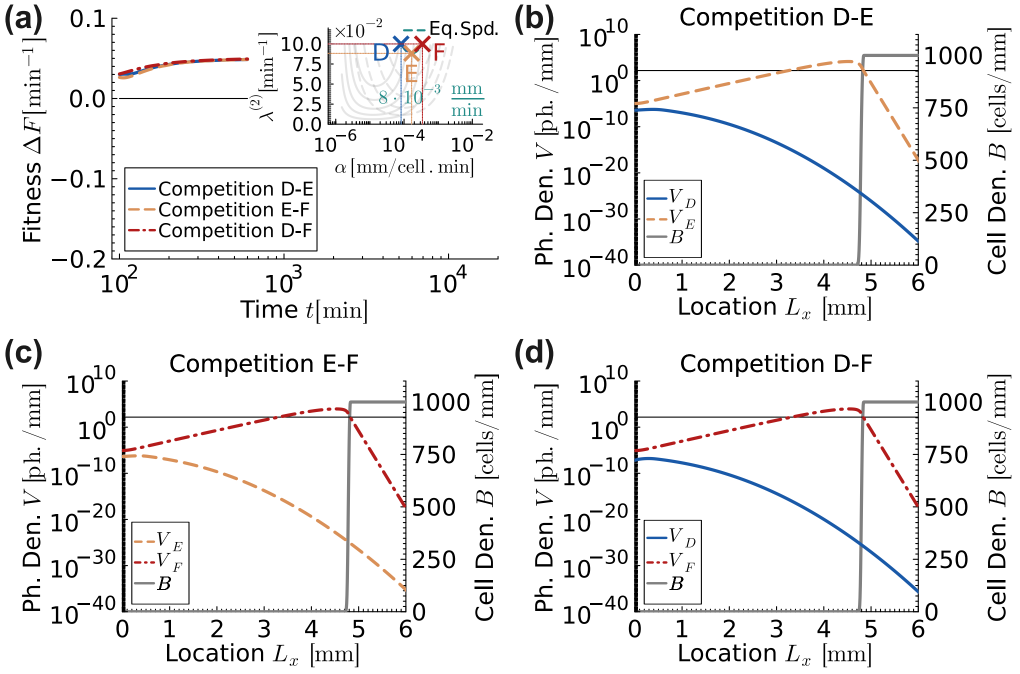


Figure B1: **Fitness and population dynamics of phages, having equal steady state plaque front expansion speeds of $8 \times 10^{-3}$ mm/min in isolation, competing in one-dimensional uniform bacterial lawn for the two phage strains model with superinfection and discreteness threshold**. (a) Relative phage fitness versus time plot with inset plot showing the three competing phages labelled as $D$, $E$, and $F$. Positive values of $\Delta F$ indicate that the second phage in the competition (E for D-E, F for E-F and F for D-F, respectively) is winning. Phages with higher adsorption rates outperform in all three competitions as the fitness lines approach positive steady state values. (b) to (d) Viral and host population distributions at time 600 minutes. Gray solid lines represent the bacterial cell densities. The thin horizontal black line marks the discreteness threshold for phage density, $\theta = 150$ ph./mm. Below this line, phages are effectively absent. Adsorption rates and second phase infection progression rates of the competing phages are mentioned in the table F1, and all remaining parameters are mentioned in table 1.

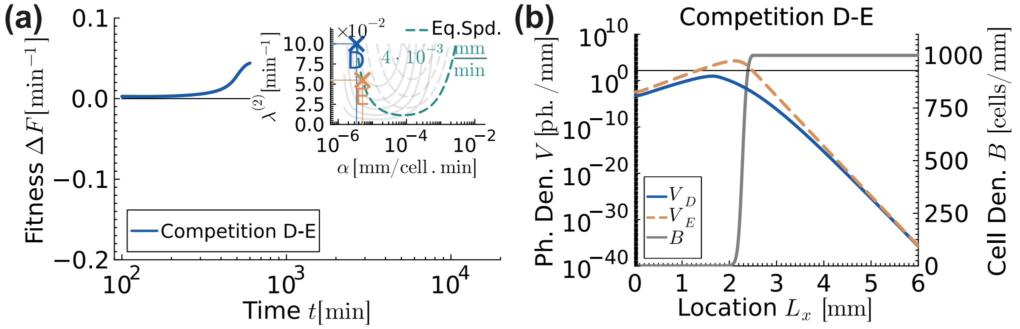


Figure B2: **Fitness and population dynamics of phages, having equal steady state plaque front expansion speeds of $4 \times 10^{-3}$ mm/min in isolation, competing in one-dimensional uniform bacterial lawn for the two phage strains model with superinfection and discreteness threshold**. (a) Relative phage fitness versus time plot with inset plot showing the two competing phages labelled as $D$ and $E$. Positive values of $\Delta F$ indicate that the second phage $E$ in the competition D-E is winning. Phage $E$ having the higher adsorption rate outperforms in competitions D-E as the fitness lines approach positive steady state values. (b) Viral and host population distributions at time 600 minutes. Gray solid lines represent the bacterial cell densities. The thin horizontal black line marks the discreteness threshold for phage density, $\theta = 150$ ph./mm. Below this line, phages are effectively absent. Adsorption rates and second phase infection progression rates of the competing phages are mentioned in the table F1, and all remaining parameters are mentioned in table 1.

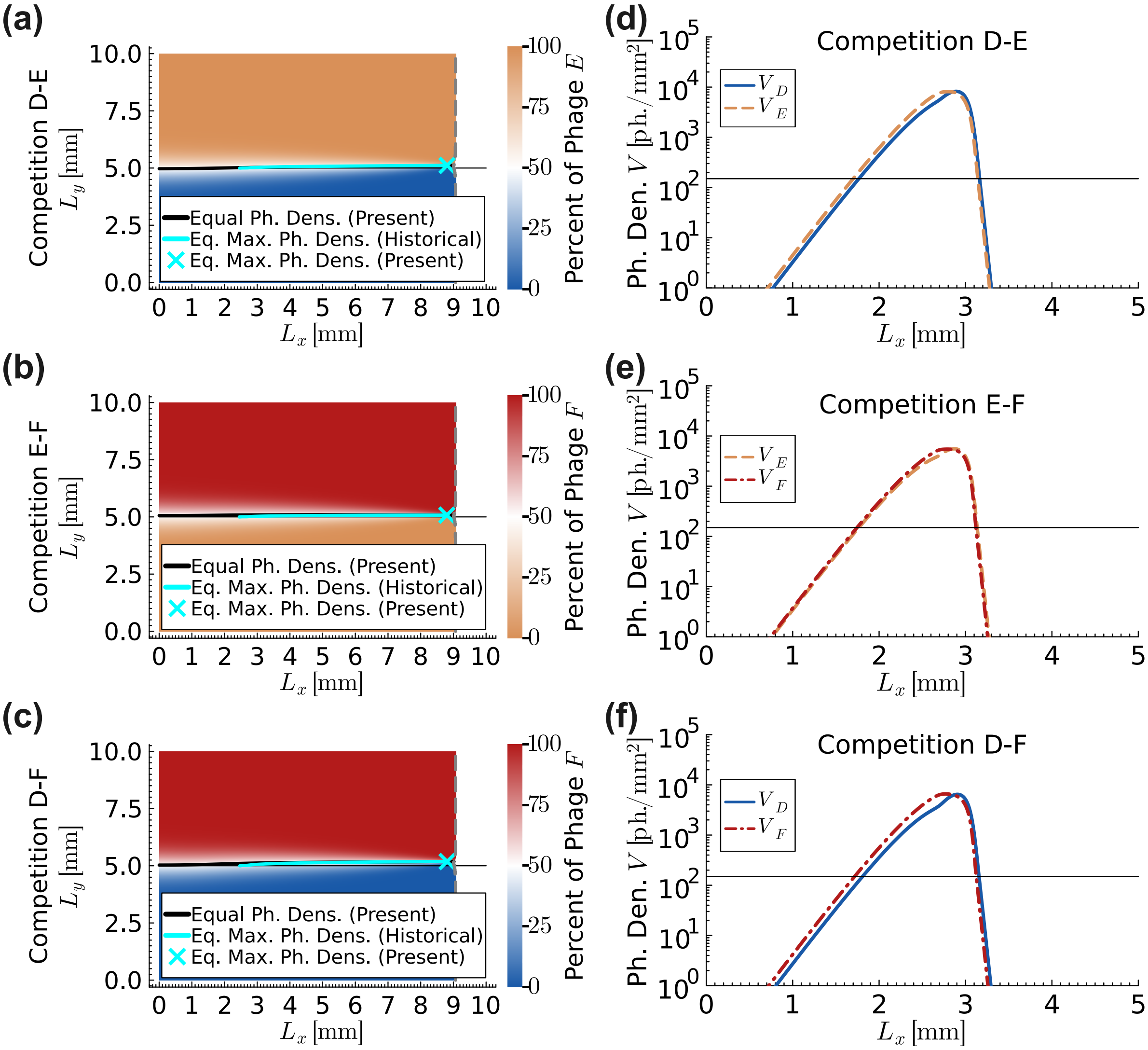


Figure B3: **Population dynamics of phages competing in a two-dimensional uniform bacterial lawn at a short timescale for the phages with equal steady state plaque front expansion speeds of $8 \times 10^{-3}$ mm/min in isolation for the model with superinfection and discreteness threshold**. Phages $D$, $E$ and $F$, having equal steady state plaque front expansion speeds of $8 \times 10^{-3}$ mm/min in isolation, are competed pair-wise in two-dimensional uniform bacterial lawn. (a) to (c) Snapshots of the simulations are given at time 800 minutes. Different phage dominated regions are shown by their respective colour labels (blue colour for phage $D$, orange colour for phage $E$ and red colour for phage $F$). Vertical gray dashed line is the plaque front. Black solid line is the instantaneous equal phage density line and cyan solid line is the historical equal phage density line. Thin horizontal black line at the centre divides the system into top and bottom halves. In all three competitions, the historical equal phage density lines (cyan) appear to be flatten over time. Phages *appear* to be neutral at short timescale once *near* steady state conditions are attained at the interface between the respective phage dominated regions. This is a universal rule that is observed in all two-dimensional competitions for all models in this research provided that the competing phages have equal steady state plaque front expansion speeds. (d) to (f) show phage density profiles in competition along the expansion axis ($x$-coordinate), evaluated at the point where the combined density of the two competing phages is maximal on the equal phage density line (i.e., cyan cross in left column) at time 50 minutes. The thin horizontal black line marks the discreteness threshold for phage density, $\theta = 150$ ph./mm. Below this line, phages are effectively absent. In all competitions, the phage having the lower adsorption rate is ahead or very close to the phage having the higher adsorption rate. Adsorption rates and second phase infection progression rates of competing phages are mentioned in the table F1, and all remaining parameters are mentioned in table 1.

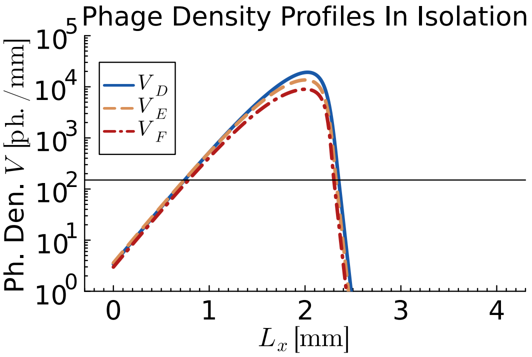


Figure B4: **Steady state initial phage density profiles after aligning the plaque fronts for the phages with equal steady state plaque front expansion speeds of $8 \times 10^{-3}$ mm/min in isolation for the model with superinfection and discreteness threshold**. Phages $D$, $E$ and $F$, having equal steady state plaque front expansion speeds of $8 \times 10^{-3}$ mm/min in isolation are allowed to expand in a one-dimensional uniform bacterial lawn in isolation until steady state is achieved. The density profiles are then aligned by defining the location of the plaque front as the position of the maximum density of total infected cells. These one-dimensional phage and cell density profiles are repeated along $y$-direction to obtain the initial condition data for the two-dimensional competitions in figure B3. Different phage densities are shown by their respective colour lines (blue solid line for phage $D$, orange dash line for phage $E$ and red dotted line for phage $F$). The figure shows that phage $D$ (*i.e.* the phage with lowest adsorption rate) leads the expansion front, followed by phage $E$, while phage $F$ (*i.e.* the phage with highest adsorption rate) lags behind. As a result, phage with a lower adsorption rate in two-dimensional competitions win at short timescales. Adsorption rates and second phase infection progression rates of phages are mentioned in the table F1, and all remaining parameters are mentioned in table 1.

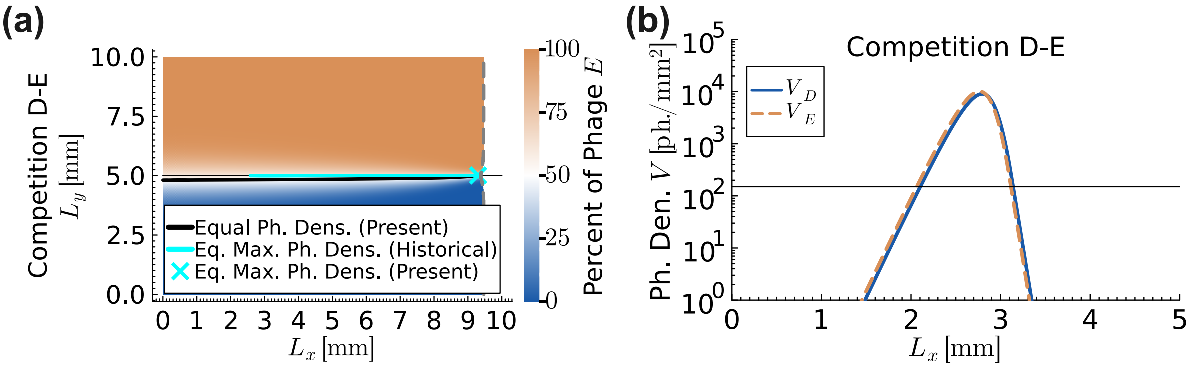


Figure B5: **Population dynamics of phages competing in a two-dimensional uniform bacterial lawn at a short timescale for the phages with equal steady state plaque front expansion speeds of $4 \times 10^{-3}$ mm/min in isolation for the model with superinfection and discreteness threshold**. Phages $D$ and $E$, having equal steady state plaque front expansion speeds of $4 \times 10^{-3}$ mm/min in isolation, are competed in two-dimensional uniform bacterial lawn. (a) Snapshot of the simulation is given at times 1700 minutes. Different phage dominated regions are shown by their respective colour labels (blue colour for phage $D$ and orange colour for phage $E$). Vertical gray dashed line is the plaque front. Black solid line is the instantaneous equal phage density line and cyan solid line is the historical equal phage density line. Thin horizontal black line at the centre divides the system into top and bottom halves. In the competition, the historical equal phage density lines (cyan) appear to be flatten over time. Phages $D$ and $E$ *appear* to be neutral on short time scales, as their life-history parameters are very similar. (b) show phage density profiles along the expansion axis ($x$-coordinate), evaluated at the point where the combined density of the two competing phages is maximal on the equal phage density line (i.e., cyan cross in left column) at time 50 minutes. The thin horizontal black line marks the discreteness threshold for phage density, $\theta = 150$ ph./mm. Below this line, phages are effectively absent. In the competition, the phage having the lower adsorption rate is marginally ahead of the phage having the higher adsorption rate. Adsorption rates and second phase infection progression rates of competing phages are mentioned in the table F1, and all remaining parameters are mentioned in table 1.

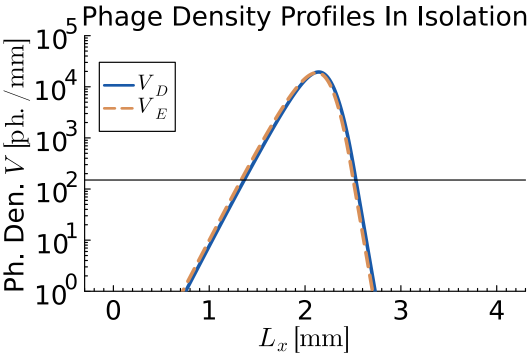


Figure B6: **Steady state initial phage density profiles after aligning the plaque fronts for the phages with equal steady state plaque front expansion speeds of $4 \times 10^{-3}$ mm/min in isolation for the model with superinfection and discreteness threshold**. Phages $D$ and $E$, having equal steady state plaque front expansion speeds of $4 \times 10^{-3}$ mm/min in isolation are allowed to expand in a one-dimensional uniform bacterial lawn in isolation until steady state is achieved. The density profiles are then aligned by defining the location of the plaque front as the position of the maximum density of total infected cells. These one-dimensional phage and cell density profiles are repeated along $y$-direction to obtain the initial condition data for the two-dimensional competitions in figure B5. Different phage densities are shown by their respective colour lines (blue solid line for phage $D$ and orange dash line for phage $E$). The figure shows that phage $D$ (*i.e.* the phage with lower adsorption rate) marginally leads the expansion front. As a result, phage with a lower adsorption rate in two-dimensional competitions win at intermediate timescales (see fig. B9). Adsorption rates and second phase infection progression rates of phages are mentioned in the table F1, and all remaining parameters are mentioned in table 1.

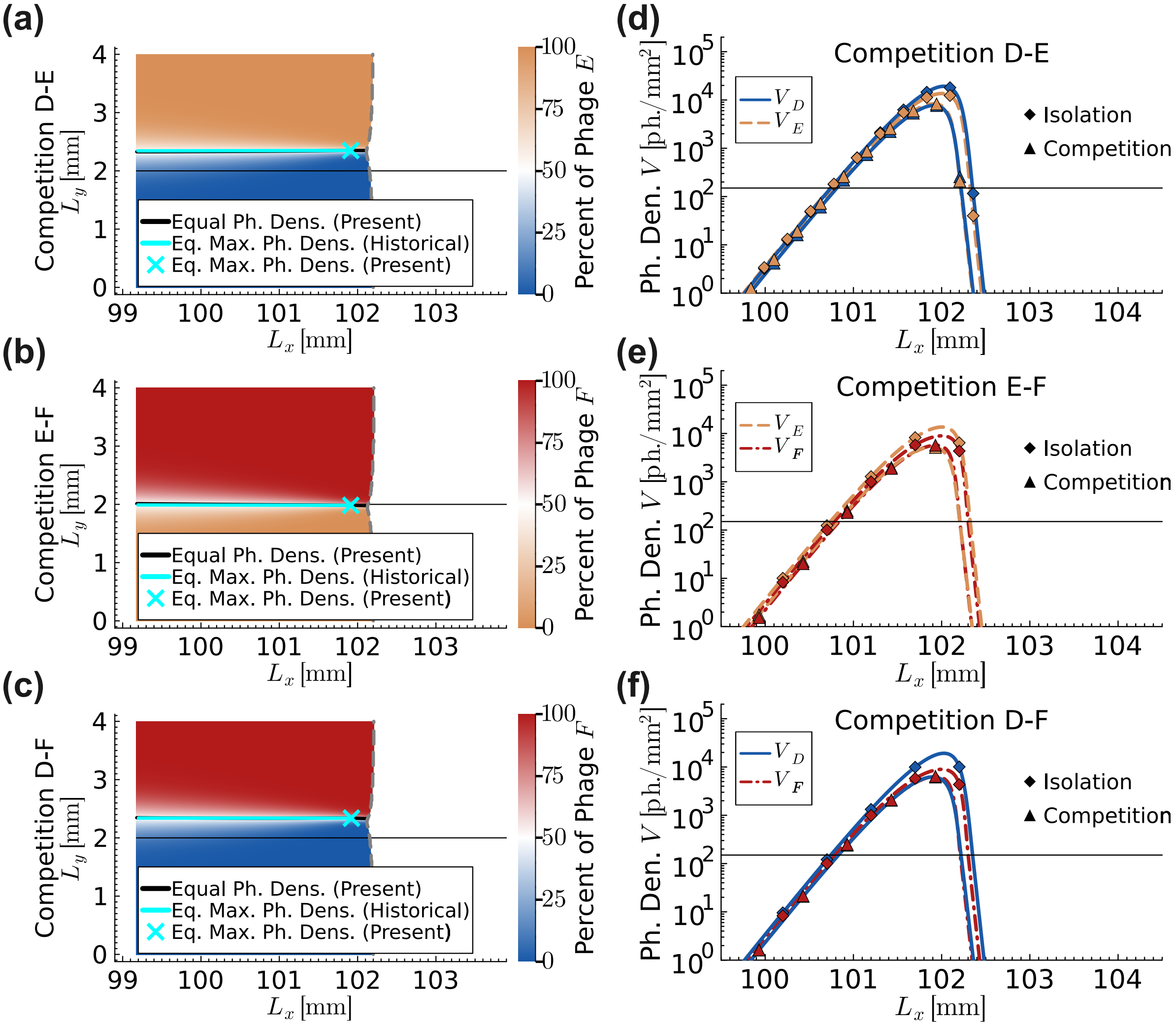


Figure B7: **Population dynamics of phages competing in a two-dimensional uniform bacterial lawn at a long timescale for the phages with equal steady state plaque front expansion speeds of $8 \times 10^{-3}$ mm/min in isolation for the model with superinfection and discreteness threshold**. Phages $D$, $E$ and $F$, having equal steady state plaque front expansion speeds of $8 \times 10^{-3}$ mm/min in isolation, are competed pair-wise in two-dimensional uniform bacterial lawn. Snapshots of the simulations are given at time 12500 minutes. Different phage dominated regions are shown by their respective colour labels (blue colour for phage $D$, orange colour for phage $E$ and red colour for phage $F$). Vertical gray dashed line is the plaque front. Black solid line is the instantaneous equal phage density line and cyan solid line is the historical equal phage density line. Thin horizontal black line at the centre divides the system into top and bottom halves. (a) to (c) In competitions D-E and D-F, the historical equal phage density lines (cyan) gradually deviated away from the thin horizontal black line over long timescale, indicating that the phages are not neutral over the long timescale. In contrast, in competition E-F, the historical equal phage density line gradually returns toward the centre of the plaque. At first glance, these competition outcomes *appear* to the trends observed in figure 10. However, this discrepancy is an artefact of the specific snapshot time chosen for the simulations (see fig. B9). (d) to (f) Curves with diamond markers show phage density profiles in isolation. Curves with triangular markers show phage density profiles in competition along the expansion axis ($x$-coordinate), evaluated at the point where the combined density of the two competing phages is maximal on the equal phage density line (i.e., cyan cross in left column). The thin horizontal black line marks the discreteness threshold for phage density, $\theta = 150$ ph./mm. Below this line, phages are effectively absent. In all competitions, the phage that attains maximum density ahead of the other phage eventually wins on the long timescale (see fig. B9). Adsorption rates and second phase infection progression rates of competing phages are mentioned in the table F1, and all remaining parameters are mentioned in table 1.

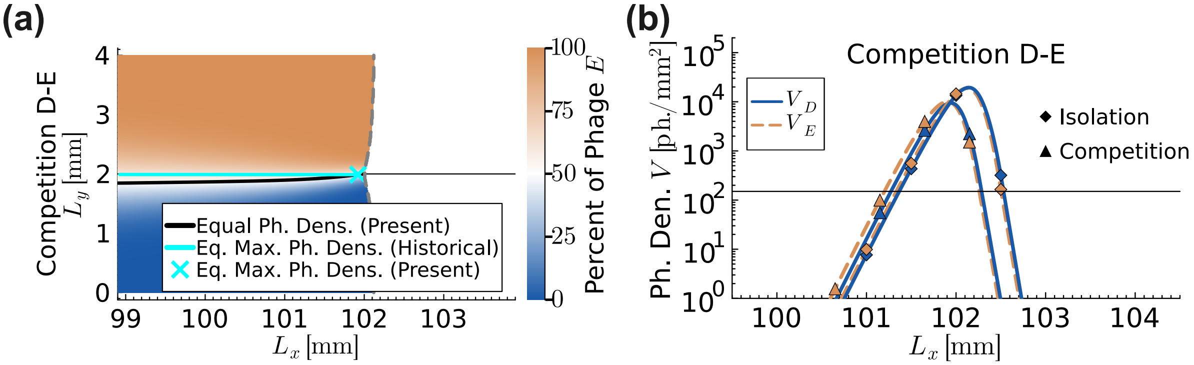


Figure B8: **Population dynamics of phages competing in a two-dimensional uniform bacterial lawn at a long timescale for the phages with equal steady state plaque front expansion speeds of $4 \times 10^{-3}$ mm/min in isolation for the model with superinfection and discreteness threshold**. Phages $D$ and $E$, having equal steady state plaque front expansion speeds of $4 \times 10^{-3}$ mm/min in isolation, are competed in two-dimensional uniform bacterial lawn. Snapshot of the simulation is given at time 25000 minutes. Different phage dominated regions are shown by their respective colour labels (blue colour for phage $D$ and orange colour for phage $E$). Vertical gray dashed line is the plaque front. Black solid line is the instantaneous equal phage density line and cyan solid line is the historical equal phage density line. Thin horizontal black line at the centre divides the system into top and bottom halves. (a) In the competition, the historical equal phage density line (cyan) remains close to the centre of the plaque even at long times, giving a misleading impression that phages $D$ and $E$ are neutral on long time scales. However, this apparent neutrality is an artefact of the specific snapshot time chosen for the simulation (see fig. B9). (b) Curves with diamond markers show phage density profiles in isolation. Curves with triangular markers show phage density profiles in competition along the expansion axis ($x$-coordinate), evaluated at the point where the combined density of the two competing phages is maximal on the equal phage density line (i.e., cyan cross in left column). The thin horizontal black line marks the discreteness threshold for phage density, $\theta = 150$ ph./mm. Below this line, phages are effectively absent. In the competition, contrary to the trends observed in figures 10 and 12, phage $E$ marginally wins at long times (see Fig. B9), even though phage $D$ attains maximum density ahead of phage $E$. This outcome likely arises because the life-history parameters of phages $D$ and $E$ are very similar and phage $E$ does not lie at the minimum of the corresponding isospeed curve (inset of fig. B2). Adsorption rates and second phase infection progression rates of competing phages are mentioned in the table F1, and all remaining parameters are mentioned in table 1.

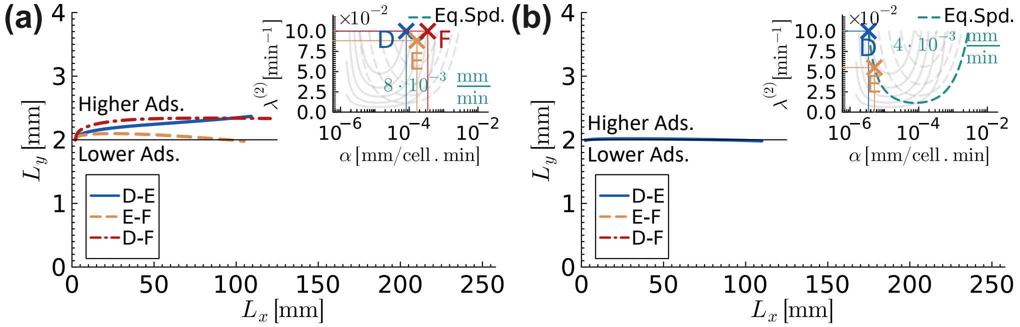


Figure B9: **Historical equal maximum phage density lines in a two-dimensional uniform bacterial lawn at a long timescale for the phages with equal steady state plaque front expansion speeds in isolation for the model with superinfection and discreteness threshold**. Simulations are initiated such that the top half of the system contained only the phage with higher adsorption rate, while the bottom half of the system contained only the phage with lower adsorption rate. The plots show that the phage with lower adsorption rate always win at short timescales, however, over long timescales the trend may shift. (a) In competition D-E, the phage $D$ wins on long timescale. In competitions E-F and D-F, phage $F$ would decisively win on long timescale if the simulations were continued further. (b) In competition D-E, phage $E$ marginally wins on long timescale. Adsorption rates and second phase infection progression rates of competing phages are mentioned in the table F1, and all remaining parameters are mentioned in table 1.

#### Appendix C. Competitions For The Model Without Superinfection But With Discreteness Threshold

To explore the effect of phage superinfection on the dynamics, competitions for the model without superinfection but with discreteness threshold are discussed in this section. Phages $D$ and $E$ are selected from isofitness curves (equal steady state plaque front expansion speeds) having the speed of $5.641 \times 10^{-3}$ mm/min.

Competition outcomes follow the same trends as observed for the model with superinfection and discreteness threshold in the main text with the only exception that instead of observing a dome-shaped behaviour of plaque speed as a function of adsorption rate, plaque speeds approach constant values at high values of adsorption rates (see figures C1 and C2).

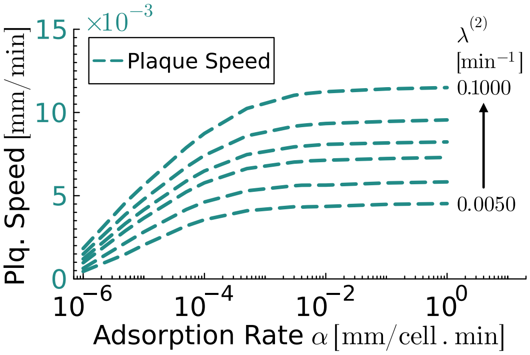


Figure C1: **Phage fitness versus adsorption rate for the one-dimensional one phage strain model without superinfection but with discreteness threshold against different values of second phase infection progression rates**. Plots for the phage fitness criteria — steady state plaque front expansion speed (dark cyan dash lines) — are shown in the figure at five different values of second phase infection progression rates: $\lambda^{(2)} = 0.005, 0.01, 0.02, 0.03, 0.05$ and $0.1$ min$^{-1}$ from bottom to top. Model parameters used to obtain the figure are mentioned in the table 1.

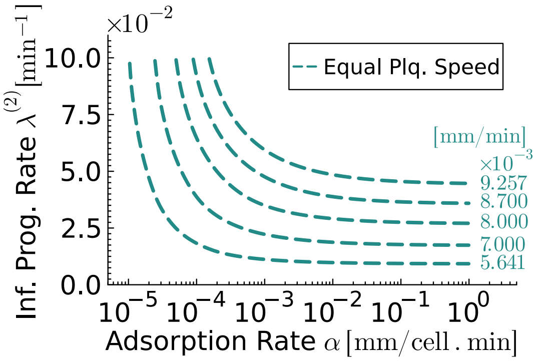


Figure C2: **Isofitness curves for the one-dimensional one phage strain model without superinfection but with discreteness threshold**. The isofitness curves provide different combinations of adsorption rates and second phase infection progression rates for phages having the same fitness levels. Isofitness curves representing the steady state plaque front expansion speeds, $5.641 \times 10^{-3}, 7 \times 10^{-3}, 8 \times 10^{-3}, 8.7 \times 10^{-3}$ and $9.257 \times 10^{-3}$ mm/min, are shown by dark cyan dash lines. Model parameters used to obtain the figure are mentioned in the table 1.

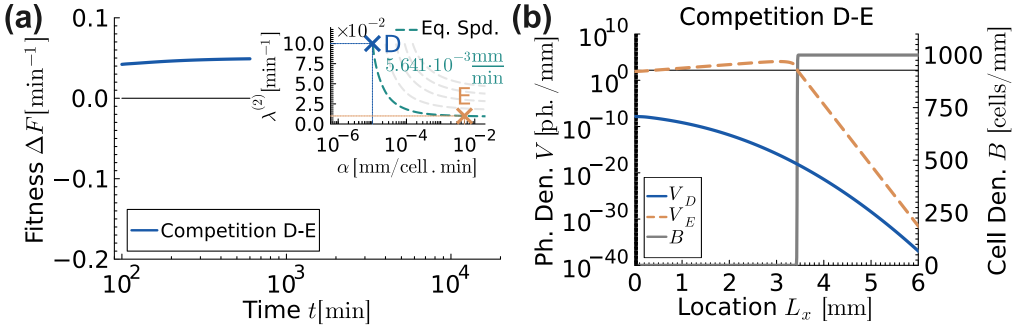


Figure C3: **Fitness and population dynamics of phages, having equal steady state plaque front expansion speeds of 5.641 $\times$ $10^{-3}$ mm/min in isolation, competing in one-dimensional uniform bacterial lawn for the two phage strains model without superinfection but with discreteness threshold**. (a) Relative phage fitness versus time plot with inset plot showing the two competing phages labelled as $D$ and $E$. Positive values of $\Delta F$ indicate that the second phage $E$ in the competition D-E is winning. Phage $E$ having the higher adsorption rate outperforms in competitions D-E as the fitness line approaches positive steady state values. (b) Viral and host population distributions at time 600 minutes. Gray solid lines represent the bacterial cell densities. The thin horizontal black line marks the discreteness threshold for phage density, $\theta = 150$ ph./mm. Below this line, phages are effectively absent. Adsorption rates and second phase infection progression rates of the competing phages are mentioned in the table F2, and all remaining parameters are mentioned in table 1.

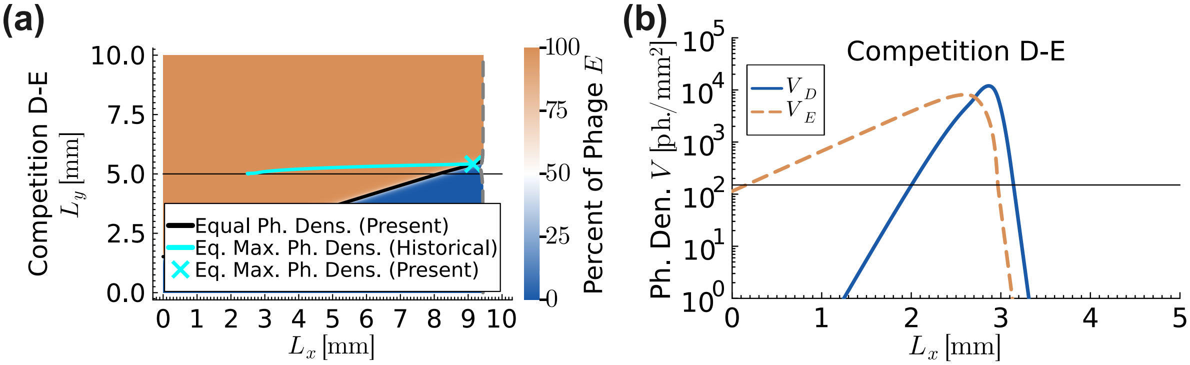


Figure C4: **Population dynamics of phages competing in a two-dimensional uniform bacterial lawn at a short timescale for the phages with equal steady state plaque front expansion speeds of 5.641 $\times$ $10^{-3}$ mm/min in isolation for the model without superinfection but with discreteness threshold**. Phages $D$ and $E$, having equal steady state plaque front expansion speeds of $5.641 \times 10^{-3}$ mm/min in isolation, are competed in two-dimensional uniform bacterial lawn. (a) Snapshot of the simulation is given at times 1200 minutes. Different phage dominated regions are shown by their respective colour labels (blue colour for phage $D$ and orange colour for phage $E$). Vertical gray dashed line is the plaque front. Black solid line is the instantaneous equal phage density line and cyan solid line is the historical equal phage density line. Thin horizontal black line at the centre divides the system into top and bottom halves. In the competition, the historical equal phage density line (cyan) appears to be flatten over time. Phages *appear* to be neutral at short timescale once *near* steady state conditions are attained at the interface between the respective phage dominated regions. This is a universal rule that is observed in all two-dimensional competitions for all models in this research provided that the competing phages have equal steady state plaque front expansion speeds. Rich dynamics are observed behind the plaque front with reference to the historical equal phage density line (cyan). In the competition, the phage $E$ dominated region is expanding into the phage $D$ dominated region below the historical equal phage density line. (b) show phage density profiles along the expansion axis ($x$-coordinate), evaluated at the point where the combined density of the two competing phages is maximal on the equal phage density line (i.e., cyan cross in left column) at time 50 minutes. The thin horizontal black line marks the discreteness threshold for phage density, $\theta = 150$ ph./mm. Below this line, phages are effectively absent. In the competition, the phage having the lower adsorption rate is ahead of the phage having the higher adsorption rate. Adsorption rates and second phase infection progression rates of competing phages are mentioned in the table F2, and all remaining parameters are mentioned in table 1.

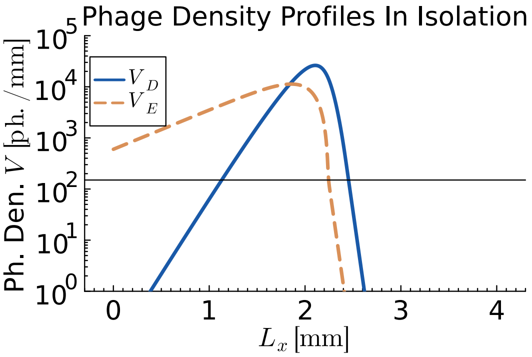


Figure C5: **Steady state initial phage density profiles after aligning the plaque fronts for the phages with equal steady state plaque front expansion speeds of 5.641 $\times$ 10$^{-3}$ mm/min in isolation for the model with superinfection and discreteness threshold**. Phages $D$ and $E$, having equal steady state plaque front expansion speeds of $5.641 \times 10^{-3}$ mm/min in isolation are allowed to expand in a one-dimensional uniform bacterial lawn in isolation until steady state is achieved. The density profiles are then aligned by defining the location of the plaque front as the position of the maximum density of total infected cells. These one-dimensional phage and cell density profiles are repeated along $y$-direction to obtain the initial condition data for the two-dimensional competitions in figure C4. Different phage densities are shown by their respective colour lines (blue solid line for phage $D$ and orange dash line for phage $E$). The figure shows that phage $D$ (*i.e.* the phage with lower adsorption rate) leads the expansion front. As a result, phage with a lower adsorption rate in two-dimensional competitions win at short timescales. Adsorption rates and second phase infection progression rates of phages are mentioned in the table F1, and all remaining parameters are mentioned in table 1.

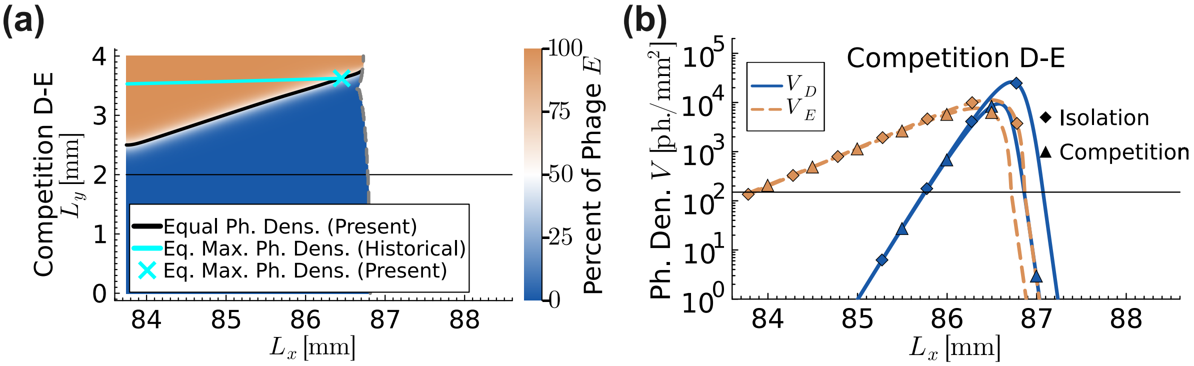


Figure C6: **Population dynamics of phages competing in a two-dimensional uniform bacterial lawn at a long timescale for the phages with equal steady state plaque front expansion speeds of 5.641 $\times$ 10$^{-3}$ mm/min in isolation for the model without superinfection but with discreteness threshold**. Phages $D$ and $E$, having equal steady state plaque front expansion speeds of $5.641 \times 10^{-3}$ mm/min in isolation, are competed in two-dimensional uniform bacterial lawn. Snapshot of the simulation is given at time 15000 minutes. Different phage dominated regions are shown by their respective colour labels (blue colour for phage $D$ and orange colour for phage $E$). Vertical gray dashed line is the plaque front. Black solid line is the instantaneous equal phage density line and cyan solid line is the historical equal phage density line. Thin horizontal black line at the centre divides the system into top and bottom halves. (a) In the competition, the historical equal phage density line (cyan) gradually deviated away from the thin horizontal black line over long timescale, indicating that the phages are not neutral over the long timescale. (b) Curves with diamond markers show phage density profiles in isolation. Curves with triangular markers show phage density profiles in competition along the expansion axis ($x$-coordinate), evaluated at the point where the combined density of the two competing phages is maximal on the equal phage density line (i.e., cyan cross in left column). The thin horizontal black line marks the discreteness threshold for phage density, $\theta = 150$ ph./mm. Below this line, phages are effectively absent. In the competition, the phage $D$ attains maximum density ahead of the other phage and eventually wins on the long timescale. Adsorption rates and second phase infection progression rates of competing phages are mentioned in the table F2, and all remaining parameters are mentioned in table 1.

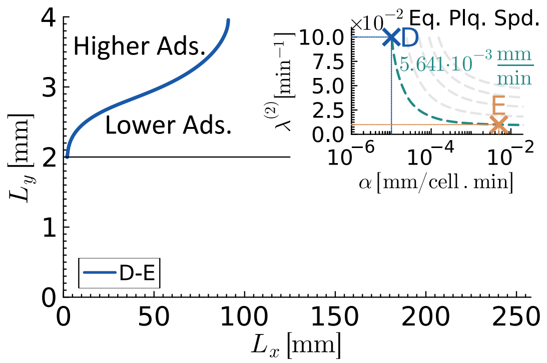


Figure C7: **Historical equal maximum phage density lines in a two-dimensional uniform bacterial lawn at a long timescale for the phages with equal steady state plaque front expansion speeds in isolation for the model without superinfection but with discreteness threshold**. Simulations are initiated such that the top half of the system contained only the phage with higher adsorption rate, while the bottom half of the system contained only the phage with lower adsorption rate. The plot shows that, in competition D-E, phage $D$ having a lower adsorption rate wins at short and long timescales. Adsorption rates and second phase infection progression rates of competing phages are mentioned in the table F2, and all remaining parameters are mentioned in table 1.

#### Appendix D. Competitions For The Model With Superinfection But Without Discreteness Threshold

To further explore the effect of phage discreteness threshold, competitions for the model with superinfection but without discreteness threshold are discussed in this section.

Phages $A$, $B$, and $C$ are selected from a single isofitness curve having equal steady state maximum phage density of 20000 ph./mm for the pair-wise competitions A-B, B-C, and A-C. Similarly, phages $D$, $E$, and $F$ are selected from a single isofitness curves having equal steady state plaque front expansion speed of $8 \times 10^{-3}$ mm/min for the pair-wise competitions D-E, E-F, and D-F. The results of this model are largely consistent with those of the model presented in the main text, except for one key difference, which is discussed below.

For this two phage strains model with superinfection but without discreteness threshold, all three phages, $D$, $E$, and $F$, having equal steady state plaque front expansion speeds in isolation, are found to be neutral in the one-dimensional pair-wise competitions in uniform bacterial lawn. Figure D4(a) is a log-log plot showing that the relative phage fitness values (from equation 7) are asymptotically approaching zero in all three competitions. Although competing phages were coexisting by the end of the simulations, phages with higher adsorption rates were found to have higher steady state maximum phage densities than the phages with lower adsorption rates (see figures D4(b) to (d)).

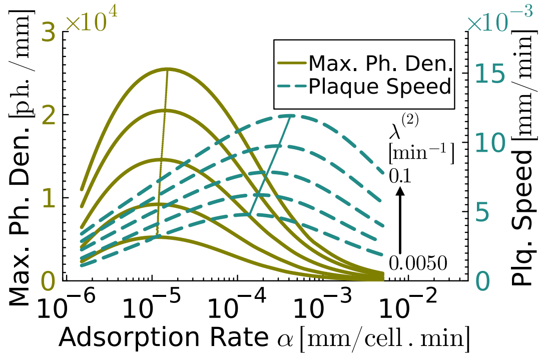


Figure D1: **Phage fitness versus adsorption rate for the one-dimensional one phage strain model with superinfection but without discreteness threshold against different values of second phase infection progression rates**. Plots for two phage fitness criteria — steady state maximum phage density at plaque front (olive solid lines) and steady state plaque front expansion speed (dark cyan dash lines) — are shown in the figure at five different values of second phase infection progression rates: $\lambda^{(2)} = 0.005, 0.01057, 0.02236, 0.04729$, and 0.1 $\text{min}^{-1}$ from bottom to top. The plots are dome-shaped, and lines tracing the peak of the domes are also shown on respective fitness plots. To obtain *near* steady state conditions, plaques are allowed to expand for a time of 1200 minutes. Asymptotic steady state values of fitness are calculated by using power-law. Model parameters used to obtain the figure are mentioned in the table 1.

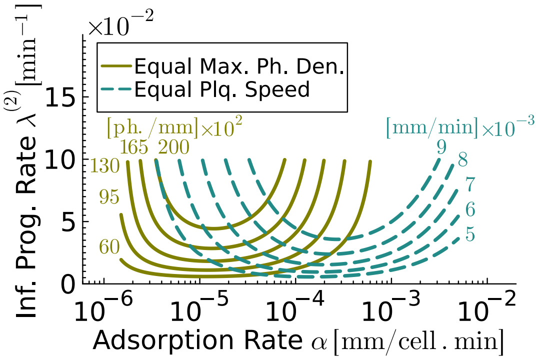


Figure D2: **Isofitness curves for the one-dimensional one phage strain model with superinfection but without discreteness threshold**. The isofitness curves provide different combinations of adsorption rates and second phase infection progression for phages having the same fitness levels. Isofitness curves representing the steady state maximum phage densities, 6000, 9500, 13000, 16500 and 20000 ph./mm, are shown by olive solid lines and isofitness curves representing the steady state plaque front expansion speeds, $5 \times 10^{-3}, 6 \times 10^{-3}, 7 \times 10^{-3}, 8 \times 10^{-3}$ and $9 \times 10^{-3}$ mm/min, are shown by dark cyan dash lines. Model parameters used to obtain the figure are mentioned in the table 1.

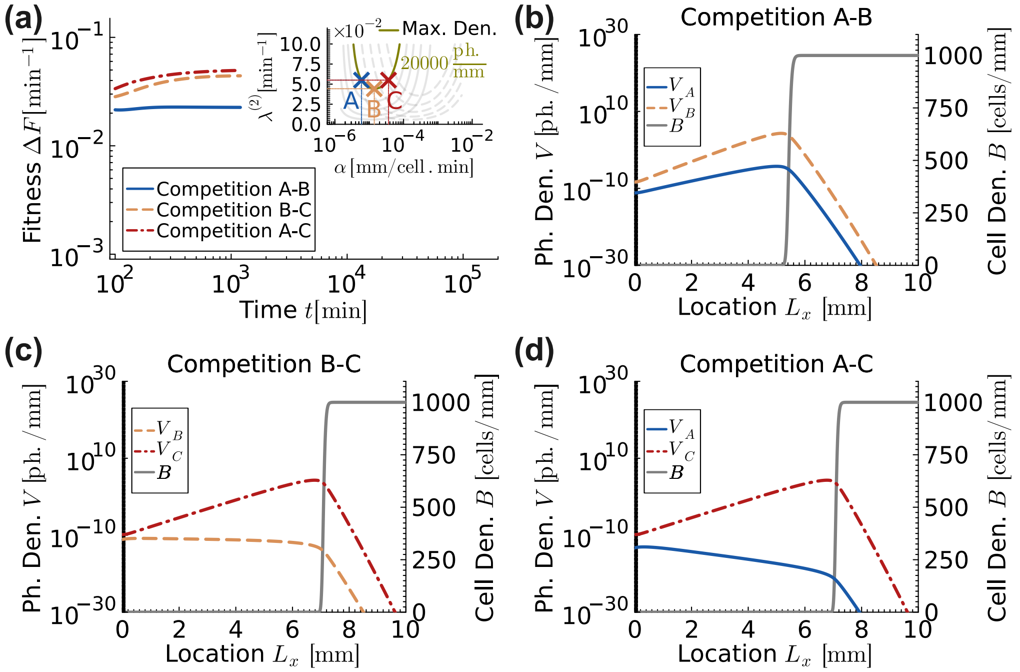


Figure D3: **Fitness and population dynamics of phages, having equal steady state maximum phage densities of 20000 ph./mm in isolation, competing in one-dimensional uniform bacterial lawn for the one phage strain model with superinfection but without discreteness threshold**. (a) Relative phage fitness versus time plot with inset plot showing the three competing phages labelled as $A$, $B$, and $C$. Positive values of $\Delta F$ indicate that the second phage in the competition (B for A-B, C for B-C and C for A-C, respectively) is winning. Phages with higher adsorption rates outperform in all three cases as the fitness lines approach positive steady state values. (b) to (d) Viral and host population distributions at time 900 minutes. Gray solid lines represent the bacterial cell densities. In all cases, the phage with higher adsorption rate eliminated the phage with lower adsorption rate. Adsorption rates and second phase infection progression rates of the competing phages are mentioned in the table F3, and all remaining parameters are mentioned in table 1.

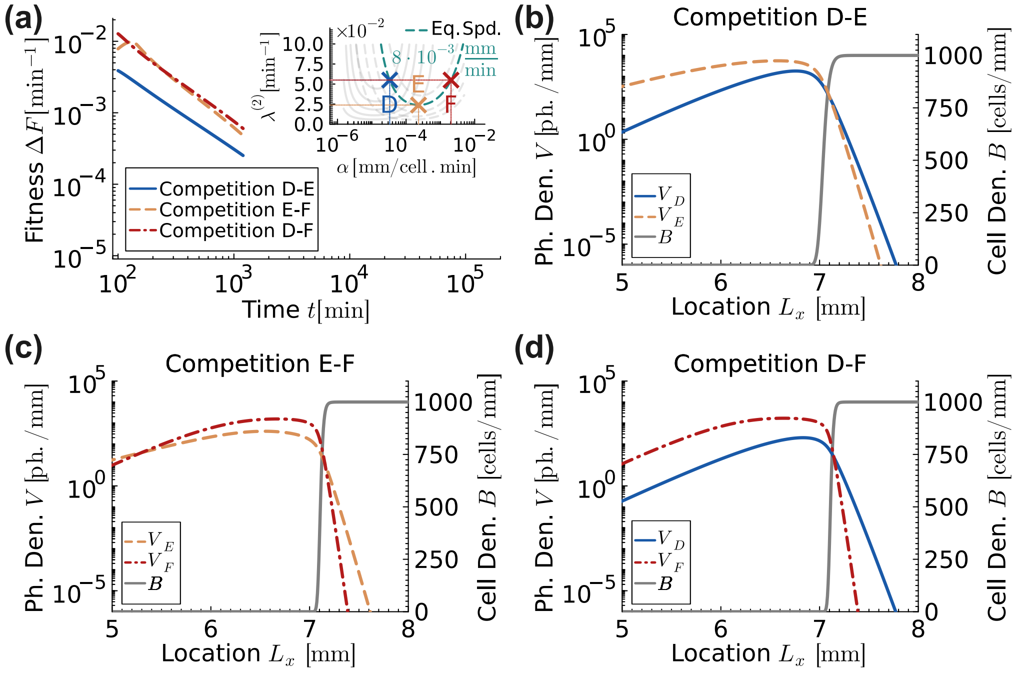


Figure D4: **Fitness and population dynamics of phages, having equal steady state plaque front expansion speeds of $8 \times 10^{-3}$ mm/min in isolation, competing in one-dimensional uniform bacterial lawn for the two phage strains model with superinfection but without discreteness threshold**. (a) Relative phage fitness versus time plot with inset plot showing the three competing phages labelled as $D$, $E$, and $F$. Positive values of $\Delta F$ indicate that the second phage in the competition (E for D-E, F for E-F and F for D-F, respectively) is winning. Competing phages are neutral in all three cases as relative phage fitness is asymptotically approaching zero in all three cases on the log-log fitness versus time plot. (b) to (d) Viral and host population distributions at time 900 minutes. Gray solid lines represent the bacterial cell densities. Competing phages continue to coexist in all three cases by the end of simulation time 900 minutes. Adsorption rates and second phase infection progression rates of the competing phages are mentioned in the table F3, and all remaining parameters are mentioned in table 1.

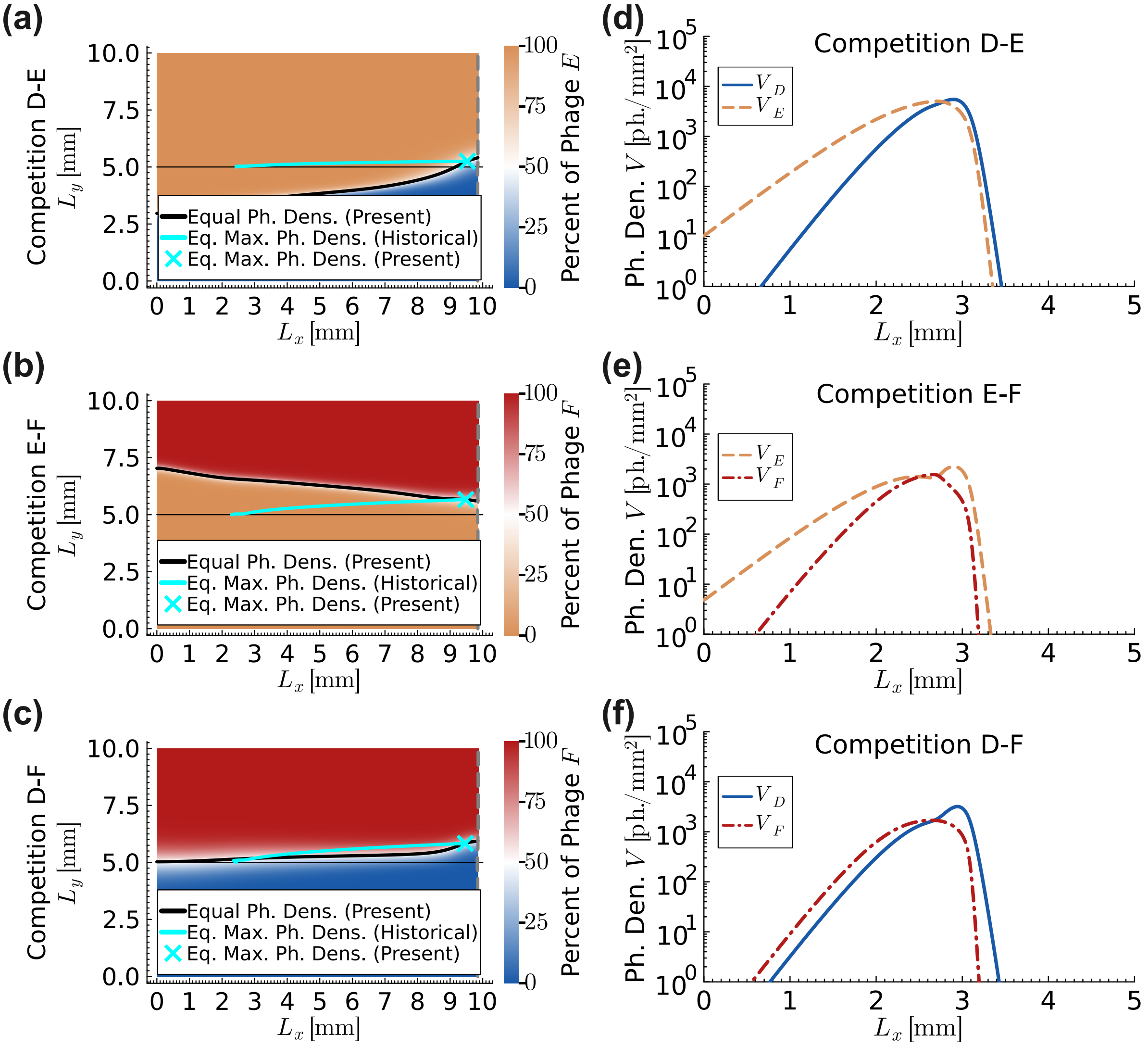


Figure D5: **Population dynamics of phages competing in a two-dimensional uniform bacterial lawn at a short timescale for the phages with equal steady state plaque front expansion speeds of $8 \times 10^{-3}$ mm/min in isolation for the model with superinfection but without discreteness threshold**. Phages $D$, $E$ and $F$, having equal steady state plaque front expansion speeds of $8 \times 10^{-3}$ mm/min in isolation, are competed pair-wise in two-dimensional uniform bacterial lawn. (a) to (c) Snapshots of the simulations are given at time 900 minutes. Different phage dominated regions are shown by their respective colour labels (blue colour for phage $D$, orange colour for phage $E$ and red colour for phage $F$). Vertical gray dashed line is the plaque front. Black solid line is the instantaneous equal phage density line and cyan solid line is the historical equal phage density line. Thin horizontal black line at the centre divides the system into top and bottom halves. In all three competitions, the historical equal phage density lines (cyan) appear to be flatten over time. Phages *appear* to be neutral at short timescale once *near* steady state conditions are attained at the interface between the respective phage dominated regions. This is a universal rule that is observed in all two-dimensional competitions for all models in this research provided that the competing phages have equal steady state plaque front expansion speeds. Rich dynamics are observed behind the plaque front with reference to the historical equal phage density line (cyan). (a) In competition D-E, the phage $E$ dominated region is expanding into the phage $D$ dominated region below the historical equal phage density line. (b) In competition E-F, the phage $E$ dominated region is expanding into the phage $F$ dominated region above the historical equal phage density line. (c) In competition D-F, the phage $F$ dominated region is expanding into the phage $D$ dominated region below the historical equal phage density line. (d) to (f) show phage density profiles in competition along the expansion axis ($x$-coordinate), evaluated at the point where the combined density of the two competing phages is maximal on the equal phage density line (i.e., cyan cross in left column) at time 50 minutes. In all competitions, the phage having the lower adsorption rate is ahead of the phage having the higher adsorption. Adsorption rates and second phase infection progression rates of competing phages are mentioned in the table F3, and all remaining parameters are mentioned in table 1.

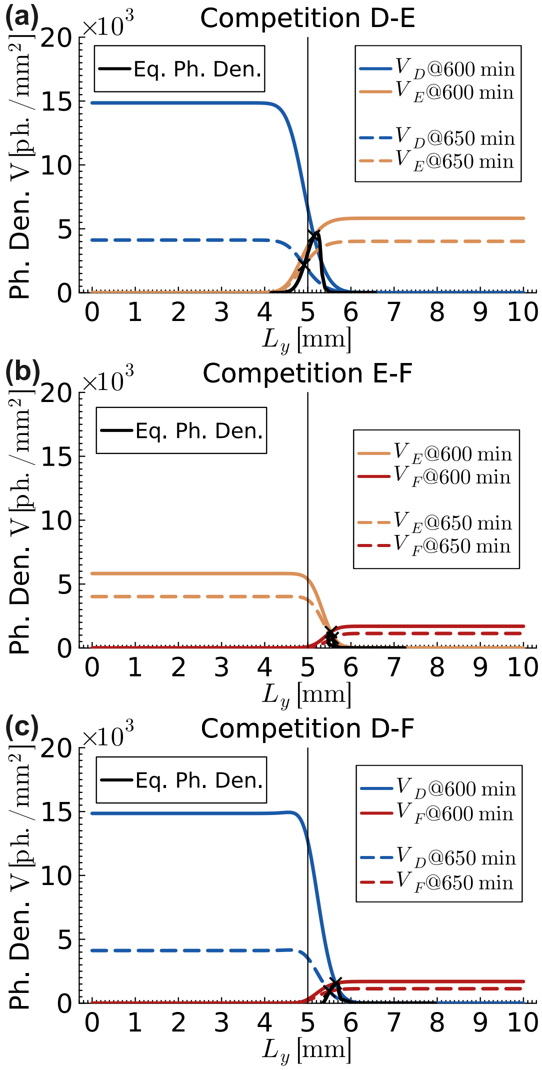


Figure D6: **Cross-sectional view of phage density profiles in a two-dimensional uniform bacterial lawn at the location $\mathbf{L_x = 7.0}$ mm for the phages with equal steady state plaque front expansion speeds in isolation for the model with superinfection but without discreteness threshold**. Phages $D$, $E$ and $F$, having equal steady state plaque front expansion speeds of $8 \times 10^{-3}$ mm/min in isolation, are competed pair-wise in two-dimensional uniform bacterial lawn. Different phage densities are shown by their respective colour labels (blue colour for phage $D$, orange colour for phage $E$ and red colour for phage $F$). Figure shows phage density profiles along $y$-coordinate at the location $L_x = 7.0$ mm at times 600 and 650 minutes as the plaque front passes through the cross section. Population densities at 600 minutes are shown by solid lines and population densities at 650 minutes are shown by dashed lines. The thin vertical black solid line at the centre divides the system into left and right halves. The thick black line traces the vertical position $L_y$ of the point on the cross section at which the two competing phages have equal densities. (a) In competition D-E, the equal population point (black cross) moved from right half to left half of the cross section because of a rapid decline in the population of phage $D$ from time 600 to 650 minutes. (b) In competition E-F, the equal population line remained on the right half of the cross section because the populations of both phages declined with similar rates. (c) In competition D-F, the equal population point (black cross) moved from right to left towards the centre of the cross section because of a rapid decline in the population of phage $D$ from time 600 to 650 minutes. However, since the population of phage $F$ was too small as compared to the population of phage $D$, the equal population line remained on the right half of the cross section at all times. Adsorption rates and second phase infection progression rates of competing phages are mentioned in the table F3, and all remaining parameters are mentioned in table 1.

From two-dimensional competitions D-E, E-F and D-F, figure D5, we found that competing phages *appears* to be neutral at the plaque front once *near* steady states were attained at the interface of respective phage dominated regions. Population dynamics is more interesting behind the plaque fronts. In competition D-E, phage $E$ dominated region expands below the historical equal phage population line and phage $D$ dominated region shrinks behind the plaque front. On the other hand, in competitions E-F, phage $E$ dominated region expands above the historical equal phage population line and phage $F$ dominated region shrinks behind the plaque front. Lastly, in competition D-F, phage $F$ dominated region slightly expands below the historical equal phage population line and phage $D$ dominated region shrinks behind the plaque front.

The population dynamics behind the plaque front is mainly dictated by the population decline rates of competing phages, which strongly depend on their infection progression rates and moderately depend on their adsorption rates. Also note that the decay rate of *free* phages is identical, $\delta = 0.05$ min$^{-1}$, for all phages. Therefore, it alone cannot provide a fitness (dis)advantage to any of the competing phages. Figure D6 shows snapshots of population distributions of phages across cross-sections at $L_x = 7.0$ mm of the two-dimensional simulations at time 600 and 650 minutes and figure D7 shows population decline rate versus time plot. In the simulations, phage $D$ experiences the highest population decline rate because it has the fastest second phase infection progression rate and the lowest adsorption rate. As a result, at any given time, it has the highest number of *free* phages in the environment that are subjected to deterioration. Phage $E$ experiences the least population decline rate because it has the lowest second phase infection progression rate and moderate adsorption rate. Therefore, phages $E$ adsorb into host cells and remain inside the cells for a longer period. While they are inside cells, they do not deteriorate. Additionally, a slower infection progression rate leads to a long tail in the Hypoexponential distribution curve (see figure E1). Consequently, a visible number of cells lyse after an extended period, releasing new phages that replenish the pool of deteriorating free phages. As a result, the overall population decline rate of phages $E$ is reduced significantly. Phage $F$ experiences a moderate population decline rate because it has the highest adsorption rate and the highest second-phase infection progression rate. Consequently, they infect cells rapidly before they deteriorate in the environment. However, they quickly lyse cells, release into the environment and experience deterioration.

Consequently, in competition D-E, at time 600 minutes, the density of phage $E$ is about one-third of the density of phage $D$ and the equal population point (black cross) is located on the right half but relatively close to the centre of the cross section. However, the population of phage $D$ declines too rapidly as compared to the population of phage $E$ from time 600 to 650 minutes. Consequently, the equal population point moved from right half to the left half of the cross section. Therefore, the phage $E$ dominated region expands while phage $D$ dominated region shrinks with time.

In competition D-F, the population distribution of phage $D$ is very similar to that observed in competition D-E. However, at the time of 600 minutes, the density of phage $F$ is just one-fifteenth of the density of phage $D$ and the equal population point (black cross) is located on the right half and further away from the centre of the cross section. Again, the population of phage $D$ declines too rapidly as compared to the population of phage $F$ from time 600 to 650 minutes. Consequently, the equal population point moved from right to left towards the centre of the cross section. Therefore, the phage $F$ dominated region expands while phage $D$ dominated region shrinks with time. However, the population of phage $F$ was too small at time 600 minutes, therefore, the equal population line (black solid line) remained on the right half of the cross section.

In case of competition E-F, the population decline rates of phage $E$ and phage $F$ are similar (also see the figure D7 in which during the period of 600 to 650 minutes, just after the plaque front passes through the cross section, the population decline rates of both phages are similar). Since the density of phage $F$ is one-fourth of the density of phage $E$ at time 600 minutes, the equal population point (black cross) is located on the right half of the cross section. Furthermore, the equal population line (black solid line) remained on the right half of the cross section because the population decline rates of both phages are similar.

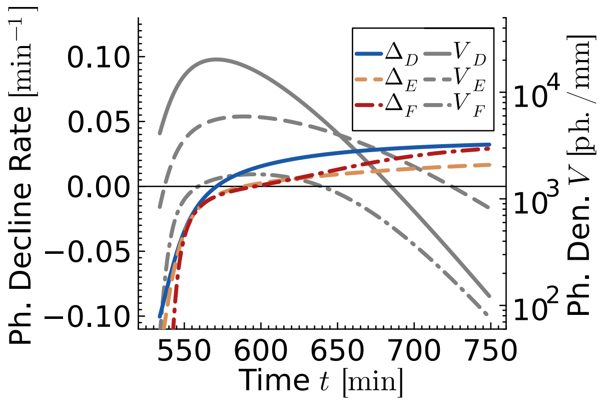


Figure D7: **Phage population decline rates of phages having equal steady state plaque front expansion speeds in one-dimensional uniform bacterial lawn for the one phage strain model with superinfection but without discreteness threshold**. Left $y$-axis is for the phage population decline rates calculated as $\Delta = -\frac{1}{V}\frac{\partial V}{\partial t}$. The negative values of decline rates imply phage growth. Right $y$-axis is for the phage density on a log-scale. The population decline rate of phage $D$ is the highest at all times. The population decline rate of phages $E$ and $F$ are similar from time 600 to 650 minutes. Adsorption rates and second phase infection progression rates of the phages are mentioned in the table F3, and all remaining parameters are mentioned in table 1.

#### Appendix E. Hypoexponential Distribution

To account for the experimentally observed variability in phage lysis times, studied both *in vitro* and *in silico* [1–4], our models assume that lysis times follow a hypoexponential distribution (also called generalised Erlang distribution).

Hypoexponential is a continuous probability distribution. It is a series of $i$ independent exponential distributions with their own rates $\lambda_i$. The mean lysis time of the phages following the $i$ stage hypoexponential distribution is given as:

$$\langle \tau \rangle = \frac{1}{\lambda^{(1)}} + \frac{1}{\lambda^{(2)}} + \cdots + \frac{1}{\lambda^{(i)}} \qquad \text{(E.1)}$$

Figure E1 shows two-stage hypoexponential distributions of phage lysis times for different values of the second phase of infection progression rates.

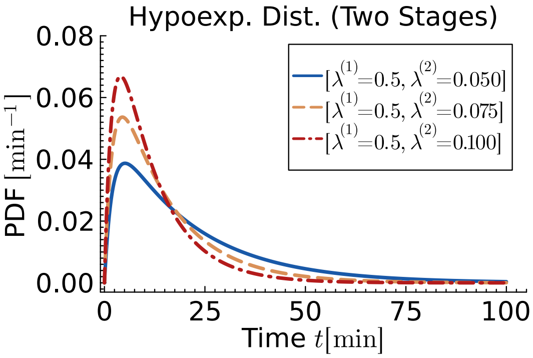


Figure E1: **Two stage hypoexponential distribution curves**. The curves show two stage hypoexponential distribution of lysis time against different values of second phase of infection progression rate.

#### Appendix F. Supplementary Tables

Table F1: **Adsorption rates and second phase infection progression rates of phages used for competition simulations for the model with superinfection and discreteness threshold**. Remaining parameters are the same for all phages and are mentioned in table 1.

| Equal Fitness Criteria | Assigned Phage Labels | Adsorption Rates $\alpha$[mm/cell · min] | Second Phase Inf. Prog. Rates $\lambda^{(2)}$[$\mathrm{min}^{-1}$] |
|---|---|---|---|
| Maximum Phage Density 16500 ph./mm | A | $3.417 \times 10^{-6}$ | 0.05500 |
| | B | $1.369 \times 10^{-5}$ | 0.03020 |
| | C | $7.240 \times 10^{-5}$ | 0.05500 |
| Plaque Front Speed $5 \times 10^{-3}$ mm/min | D | $1.132 \times 10^{-5}$ | 0.05500 |
| | E | $9.568 \times 10^{-5}$ | 0.02028 |
| | F | $8.997 \times 10^{-4}$ | 0.05500 |
| | U | $3.588 \times 10^{-5}$ | 0.02510 |
| Plaque Front Speed $4.4862 \times 10^{-3}$ mm/min | W | $9.568 \times 10^{-5}$ | 0.01522 |
| Plaque Front Speed $8 \times 10^{-3}$ mm/min | D | $8.190 \times 10^{-5}$ | 0.10000 |
| | E | $1.669 \times 10^{-4}$ | 0.08827 |
| | F | $3.460 \times 10^{-4}$ | 0.10000 |
| Plq. Front Speed $4 \times 10^{-3}$ mm/min | D | $3.452 \times 10^{-6}$ | 0.10000 |
| | E | $5.164 \times 10^{-6}$ | 0.05500 |

Table F2: **Adsorption rates and second phase infection progression rates of phages used for competition simulations for the model without superinfection but with discreteness threshold**. Remaining parameters are the same for all phages and are mentioned in table 1.

| Equal Fitness Criteria | Assigned Phage Labels | Adsorption Rates $\alpha$[mm/cell · min] | Second Phase Inf. Prog. Rates $\lambda^{(2)}$[$\mathrm{min}^{-1}$] |
|---|---|---|---|
| Plq. Front Speed $5.6412 \times 10^{-3}$ mm/min | D | $1.009 \times 10^{-5}$ | 0.10000 |
| | E | $5.000 \times 10^{-3}$ | 0.01000 |

Table F3: **Adsorption rates and second phase infection progression rates of phages used for competition simulations for the model with superinfection but without discreteness threshold**. Phages $A$, $B$, and $C$ have the same steady state maximum phage density of 20000 ph./mm whereas phages $D$, $E$, and $F$ have the same steady state plaque front expansion speed of $8 \times 10^{-3}$ mm/min. Remaining parameters are the same for all phages and are mentioned in table 1.

| **Equal Fitness Criteria** | **Assigned Phage Labels** | **Adsorption Rates** $\alpha$[**mm/cell · min**] | **Second Phase Inf. Prog. Rates** $\lambda^{(2)}$[$\mathbf{min^{-1}}$] |
|---|---|---|---|
| Maximum Phage Density 20000 ph./mm | A | $5.962 \times 10^{-6}$ | 0.05500 |
| | B | $1.406 \times 10^{-5}$ | 0.04428 |
| | C | $3.655 \times 10^{-5}$ | 0.05500 |
| Plaque Front Speed $8 \times 10^{-3}$ mm/min | D | $3.335 \times 10^{-5}$ | 0.05500 |
| | E | $2.311 \times 10^{-4}$ | 0.02385 |
| | F | $2.048 \times 10^{-3}$ | 0.05500 |
